# Neurocognitive reorganization between crystallized intelligence, fluid intelligence and white matter microstructure in two age-heterogeneous developmental cohorts

**DOI:** 10.1101/593509

**Authors:** Ivan L. Simpson-Kent, Delia Fuhrmann, Joe Bathelt, Jascha Achterberg, Gesa Sophia Borgeest, the CALM Team, Rogier A. Kievit

**Affiliations:** MRC Cognition and Brain Sciences Unit, University of Cambridge, Cambridge, Cambridgeshire, CB2 7EF, UK; Dutch Autism & ADHD Research Center, Brain & Cognition, University of Amsterdam, 1018 WS Amsterdam, Netherlands

**Keywords:** Neurocognitive reorganization, Crystallized intelligence, Fluid intelligence, White matter, Structural equation modelling

## Abstract

Despite the reliability of intelligence measures in predicting important life outcomes such as educational achievement and mortality, the exact configuration and neural correlates of cognitive abilities remain poorly understood, especially in childhood and adolescence. Therefore, we sought to elucidate the factorial structure and neural substrates of child and adolescent intelligence using two cross-sectional, developmental samples (CALM: N=551 (N=165 imaging), age range: 5-18 years, NKI-Rockland: N=337 (N=65 imaging), age range: 6-18 years). In a preregistered analysis, we used structural equation modelling (SEM) to examine the neurocognitive architecture of individual differences in childhood and adolescent cognitive ability. In both samples, we found that cognitive ability in lower and typical-ability cohorts is best understood as two separable constructs, crystallized and fluid intelligence, which became more distinct across development, in line with the age differentiation hypothesis. Further analyses revealed that white matter microstructure, most prominently the superior longitudinal fasciculus, was strongly associated with crystallized (gc) and fluid (gf) abilities. Finally, we used SEM trees to demonstrate evidence for developmental reorganization of gc and gf and their white matter substrates such that the relationships among these factors dropped between 7-8 years before increasing around age 10. Together, our results suggest that shortly before puberty marks a pivotal phase of change in the neurocognitive architecture of intelligence.

## 1. Introduction

Intelligence measures have repeatedly been shown to predict important life outcomes such as educational achievement (Deary et al., 2007) and mortality (Calvin et al., 2011). Modern investigations of intelligence began over 100 years ago, when Spearman first proposed *g* (for ‘general intelligence’) as the underlying factor behind his positive manifold of cognitive ability and established intelligence as a central theme of psychological research (Spearman, 1904). Cattell proposed a division of Spearman’s *g*-factor into two separate yet related constructs, crystallized (gc) and fluid (gf) intelligence (Cattell, 1967). Cattell suggested that gc represents the capacity to effectively complete tasks based on acquired knowledge and experience (e.g. arithmetic, vocabulary) whereas gf refers to one’s ability to solve novel problems without task-specific knowledge, relying on abstract thinking and pattern recognition (see also Deary et al., 2010).

Although other detailed conceptualizations of intelligence are available (e.g. Schneider and McGrew, 2012) fluid and crystallized intelligence have proven especially insightful regarding developmental changes in intelligence. For instance, current understanding of lifespan trajectories of gc and gf using cross-sectional (Horn and Cattell, 1967) and longitudinal (McArdle et al., 2000; Schaie, 1994) cohorts indicates that gc slowly improves until late age while gf increases into early adulthood before steadily decreasing. However, most of the literature on individual differences between gc and gf has focused on early to late adulthood. As a result, considerably less is known about the association between gc and gf in childhood and adolescence (but see Hülür et al., 2011).

There has, however, been a recent rise in interest in this topic in child and adolescent samples. For instance, research on age-related differentiation and its inverse, age dedifferentiation, in younger samples has greatly expanded since first being pioneered in the middle of the 20th century (Garrett, 1946). According to the age differentiation hypothesis, cognitive factors become less correlated (more differentiated) with increasing age. For example, the relationship (covariance) between gc and gf would decrease as children age into adolescence, suggesting that cognitive abilities increasingly specialize into adulthood. In contrast, the age dedifferentiation hypothesis predicts that cognitive abilities become more strongly related (less differentiated) throughout development. In this case, gc and gf covariance would increase between childhood and adolescence, potentially indicating a strengthening of the *g-*factor across age. However, despite its increased attention in the literature, the debate remains unsolved as evidence in support of both hypotheses has been found (Bickley et al., 1995; de Mooij et al., 2018; Gignac, 2014; Hülür et al., 2011; Juan-Espinosa et al., 2000; Tideman and Gustafsson, 2004). Together, this literature highlights the importance of a lifespan perspective on theories of cognitive development, as neither age differentiation nor dedifferentiation may be solely able to capture the dynamic changes that occur from childhood to adolescence and (late) adulthood (Hartung et al., 2018).

The introduction of non-invasive brain imaging technology has complemented conventional psychometric approaches by allowing for fine-grained probing of the neural bases of human cognition. A particular focus in developmental cognitive neuroscience has been the study of white matter using techniques such as diffusion-weighted imaging, which allows for the estimation of white matter microstructure (Wandell, 2016). Both cross-sectional and longitudinal research in children and adolescents using fractional anisotropy (FA), a commonly used estimate of white matter integrity, have consistently revealed strong correlations between FA and cognitive ability using tests of working memory, verbal and non-verbal performance (Koenis et al., 2015; Krogsrud et al., 2018; Peters et al., 2014; Tamnes et al., 2010; Urger et al., 2015). Moreover, recent research has also found associations between the corpus callosum (Navas-Sánchez et al., 2014; Westerhausen et al., 2018) association fibers (e.g. inferior longitudinal fasciculus, see Peters et al., 2014), the superior longitudinal fasciculus (Urger et al., 2015), and differences in cognitive ability, suggesting the importance of white matter integrity across large coordinated brain networks for high cognitive performance. However, interpretations of these studies are limited due to restricted cognitive batteries (e.g. small number of tests used) and a dearth of theory-driven statistical analyses (e.g. structural equation modelling). Interestingly, preliminary evidence suggests that the associations between brain structure and cognitive performance may not be static during development. For instance, Koenis et al., 2018 observed increased correlation between cognitive performance and FA-derived metric changes over adolescence.

For these reasons, several outstanding questions in the developmental cognitive neuroscience of intelligence remain: 1) Are the white matter substrates underlying intelligence in childhood and adolescence best understood as a single global factor or do individual tracts provide specific contributions to gc and gf?, 2) If they are specific, are the tract contributions identical between gc and gf?, and 3) Does this brain-behavior mapping change in development (e.g. age differentiation/dedifferentiation or both)?

To examine these questions, our <u>preregistered</u> hypotheses are as follows:

1. gc and gf are separable constructs in childhood and adolescence. More specifically, the covariance among scores on cognitive tests are more adequately captured by the two-factor (gc-gf) model as opposed to a single-factor (e.g. *g*) model.
2. The covariance between gc and gf differs (decreases) across childhood and adolescence.
3. White matter tracts make unique complementary contributions to gc and gf.
4. The contributions of these tracts to gc and gf differ (decrease) with age.

To address these questions, we examined the relationship between gc and gf in two large cross-sectional child and adolescent samples. The first is the Centre for Attention, Learning and Memory (CALM, see Holmes et al., 2019). This sample, included in our <u>preregistration</u>, was recruited atypically (see Methods for more detail) and generally includes children with slightly lower cognitive abilities than age-matched controls. To examine whether findings from CALM would generalize to other samples, we also conducted non-preregistered analyses on the Nathan Kline Institute (NKI)-Rockland Sample, a cohort with similar population demographics to the United States (e.g. race and socioeconomic status, see Table 1 of Nooner et al., 2012). All analyses were carried out using structural equation modelling (SEM), a multivariate statistical framework combining factor and path analysis to examine the extent to which causal hypotheses concerning latent (unobserved, e.g. *g*) and manifest (observed, e.g. cognitive tests scores) variables (Schreiber et al., 2006) are in line with the observed data. Taken together, this paper sought to investigate the relationship between measures of intelligence (gc and gf) and white matter connectivity in typically and atypically (struggling learners) developing children and adolescents.

**Table 1.**
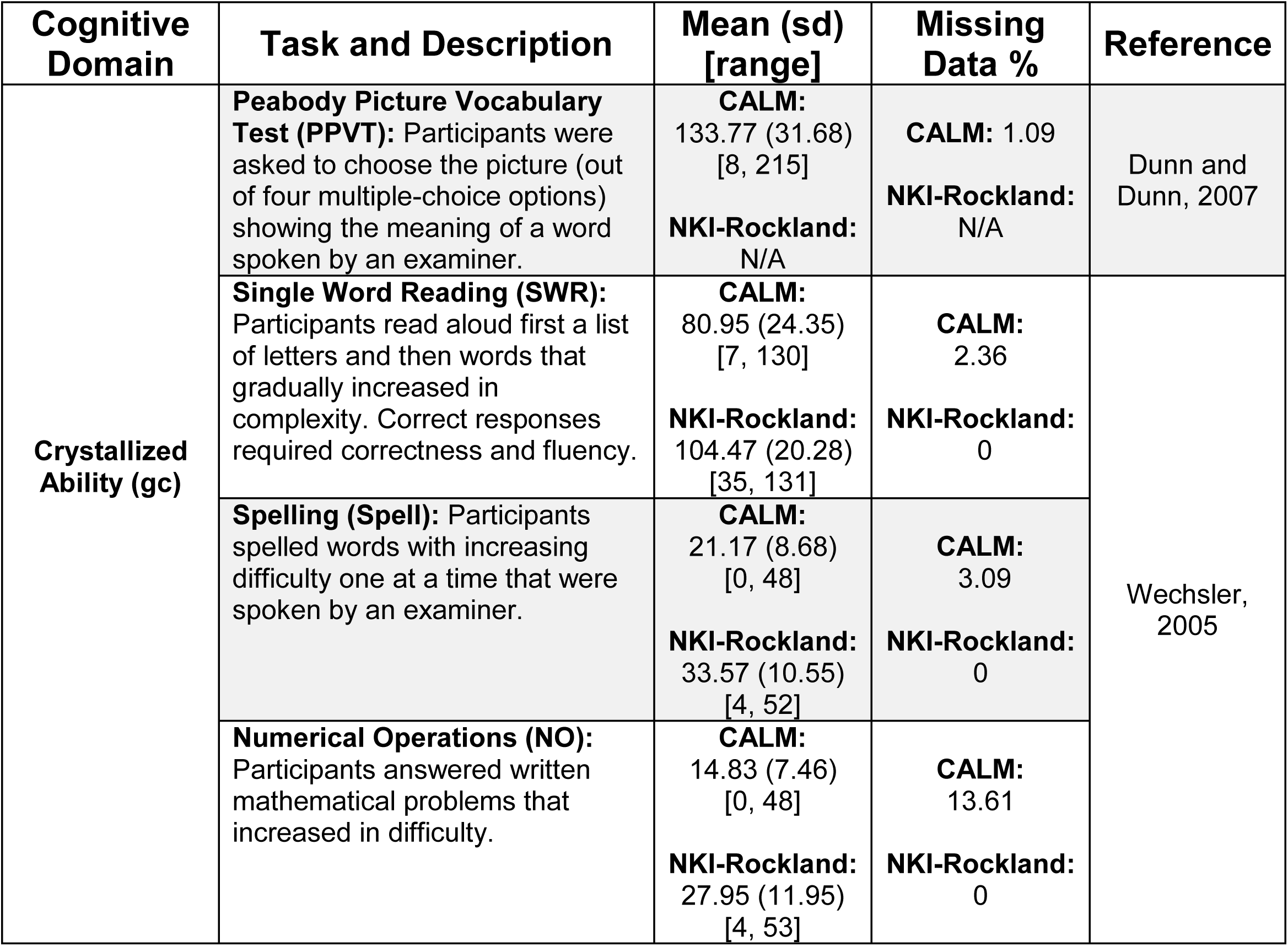
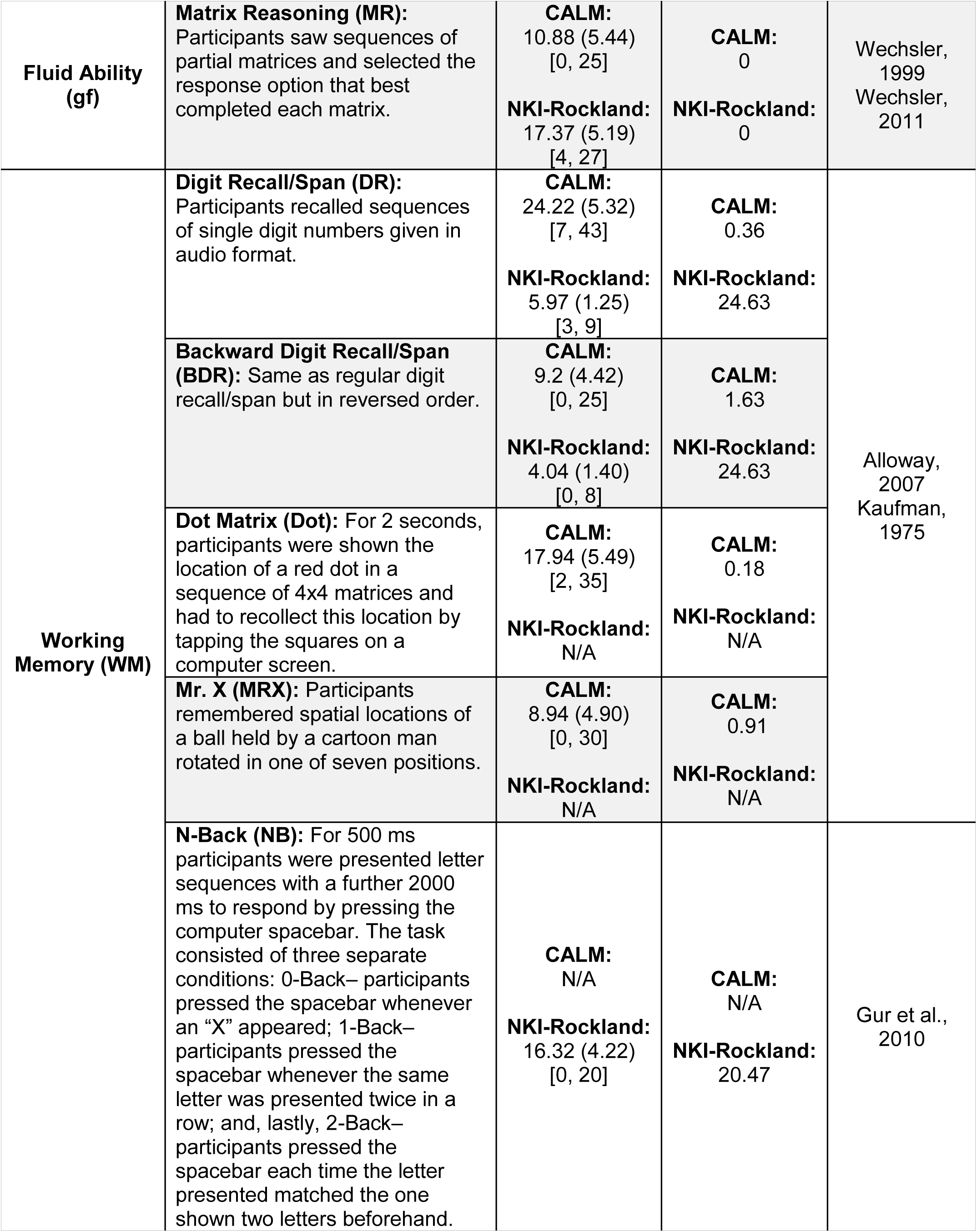
List, descriptions and summary statistics (mean, standard deviation, range, and percentage of missing data) of cognitive assessments used in CALM & NKI-Rockland samples.

## 2. Methods

### 2.1.1 Participants

For the CALM sample, we analyzed the most recent data release (N=551; 170 female, 381 male^1^; age range=5.17-17.92 years) at the time of preregistration (see https://aspredicted.org/5pz52.pdf). Participants were recruited based on referrals made for possible attention, memory, language, reading and/or mathematics problems (Holmes et al., 2019). Participants with or without formal clinical diagnosis were referred to CALM. Exclusion criteria included known significant and uncorrected problems in vision or hearing and a native language other than English. A subset of participants completed MRI scanning (N=165; 56 female, 109 male; age range=5.92-17.92 years). For more information about CALM, see http://calm.mrc-cbu.cam.ac.uk/.

Next, to assess the generalizability of our findings in CALM, we used a non-preregistered subset of the data from the Nathan Kline Institute (NKI)-Rockland Sample (cognitive data: N=337; 149 female, 188 male; age range=6.12-17.94 years; neural data: N=65; 27 female, 38 male; age range=6.97-17.8 years). This multi-institutional initiative recruited a lifespan (aged between 6 and 85 years), community-ascertained sample (Nooner et al., 2012). We chose this sample due to its representativeness (demographics resemble those of the United States population) and the fact that its cognitive battery assessments closely-matched CALM. For more information about the NKI-Rockland Sample and its procedures, see http://rocklandsample.org/. Also see Fig. 1 for age distributions of CALM and NKI-Rockland. These same two cohorts were used in a recent paper to address a distinct set of questions (Fuhrmann et al., 2019).

**Fig. 1.**
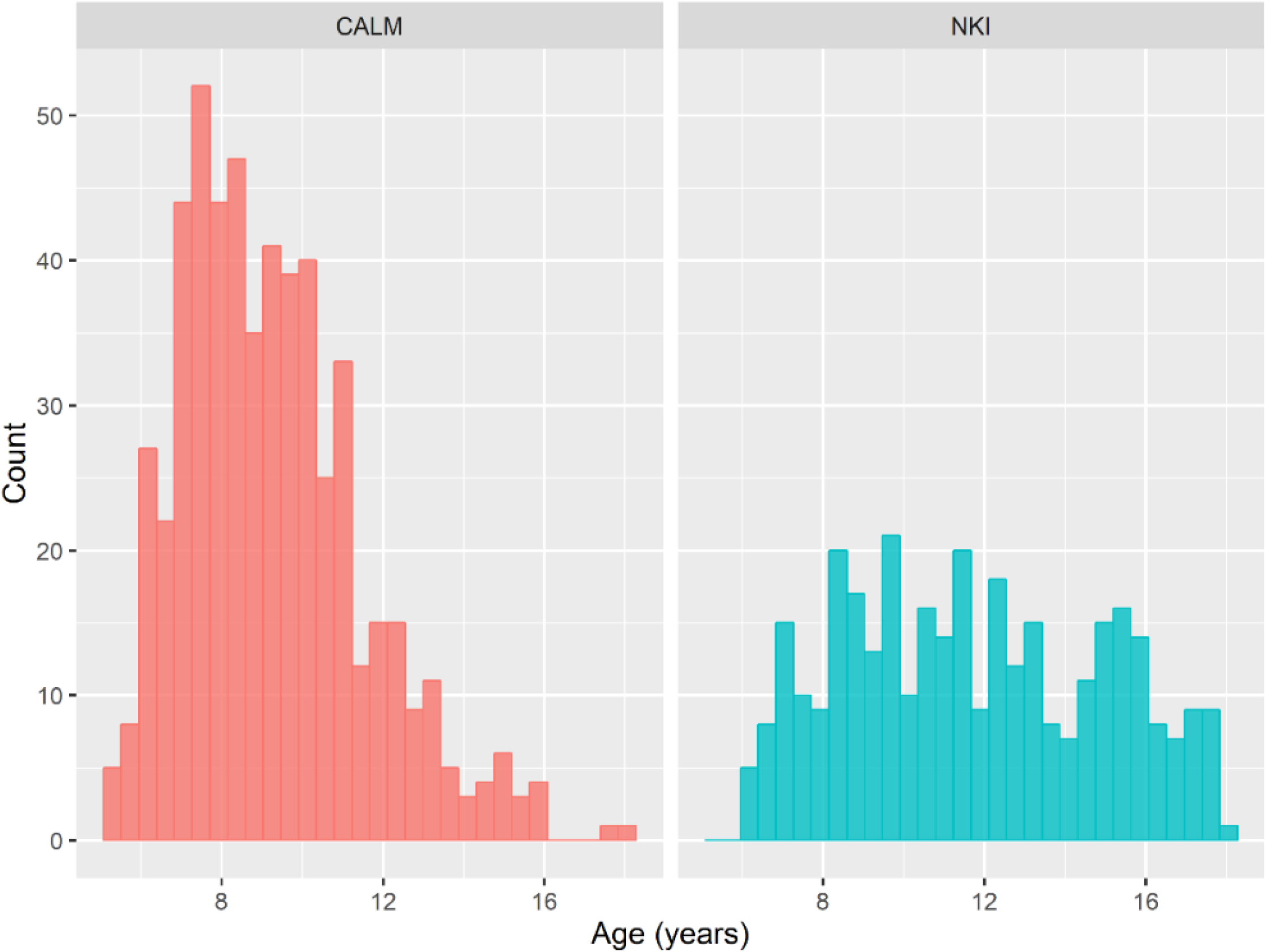
Histograms of age distributions for CALM and NKI-Rockland samples.

### 2.1.2 Cognitive assessments: gc, gf, and working memory

All cognitive data from the CALM sample were collected on a one-to-one basis by an examiner in a dedicated child-friendly testing room. The test battery included a wide range of standardized assessments of cognition and learning (Holmes et al., 2019). Participants were given regular breaks throughout the session. Testing was divided into two sessions for participants who struggled to complete the assessments in one sitting. For analyses of the NKI-Rockland Sample cohort, we matched tasks used in CALM except for the Peabody Picture Vocabulary Test, Dot Matrix, and Mr. X, which were only available for CALM. For the NKI-Rockland Sample, we included the N-Back task, which is not available in CALM (Nooner et al., 2012). In both samples, only raw scores obtained from assessments were included in analyses. Due to varying delays between recruitment and testing in the NKI-Rockland cohort, we only used cognitive test scores completed no later than six months after initial recruitment. The cognitive tasks are further described in Table 1; the raw scores are depicted in Fig. 2.

**Fig. 2.**
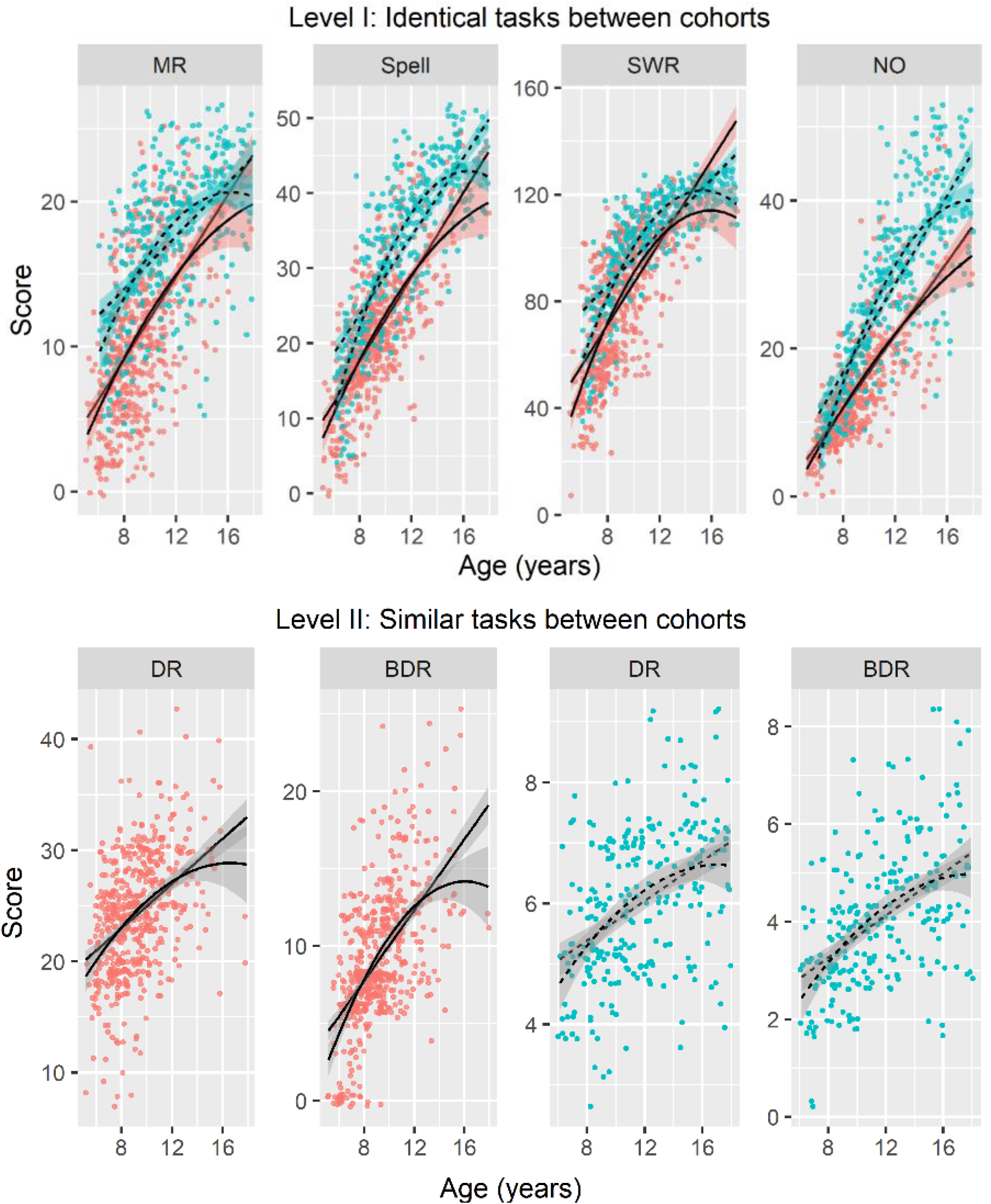
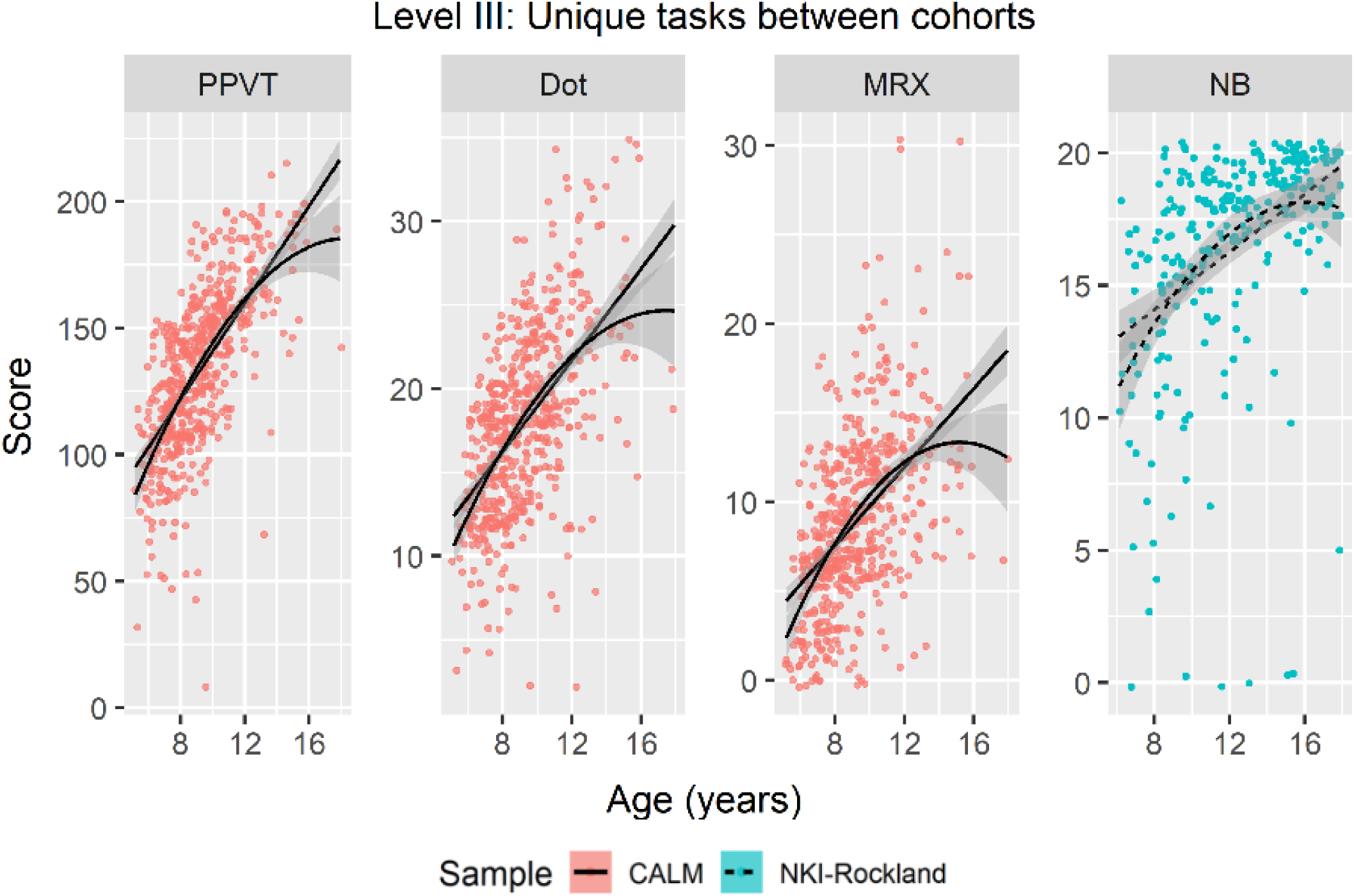
Scatterplots of cognitive task scores across age for CALM and NKI-Rockland samples. Lines and shades reflect linear and polynomial fit and 95% confidence intervals, respectively. Solid lines: CALM. Dashed lines: NKI-Rockland. Abbreviations: matrix reasoning (MR), spelling (Spell), single word reading (SWR), numerical operations (NO), forward digit recall/span (DR), backward digit recall/span (BDR), Peabody Picture Vocabulary Test (PPVT), Dot Matrix (Dot), Mr. X (MRX), N-Back (NB).

The working memory tasks, while measuring the same cognitive abilities, used different test batteries and scoring protocols. For CALM, working memory scores indicate the total number of correctly recalled digits across all trials while the NKI-Rockland scores were transformed into a span score. Due to this discrepancy (see References in Table 1 and Alloway et al., 2008 for statistical comparisons between the batteries), these tasks are plotted separately (see Fig. 2).

### 2.1.3 MRI acquisition

The CALM sample neuroimaging data were obtained at the MRC Cognition and Brain Sciences Unit, Cambridge, UK. Scans were acquired on the Siemens 3 T Tim Trio system (Siemens Healthcare, Erlangen, Germany) via 32-channel quadrature head coil. All T1-weighted volume scans were acquired using a whole brain coverage 3D magnetization-prepared rapid acquisition gradient echo (MPRAGE) sequence with 1mm isotropic image resolution with the following parameters: Repetition Time (TR)=2250 milliseconds; Echo Time (TE)=3.02 milliseconds; Inversion Time (TI)=900 milliseconds; flip angle=9 degrees; voxel dimensions=1 mm isotropic; GRAPPA acceleration factor=2. Diffusion-Weighted Images (DWI) were acquired using a Diffusion Tensor Imaging (DTI) sequence with 64 diffusion gradient directions with a b-value of 1000 s/mm2, plus one image acquired with a b-value of 0. Other relevant parameters include: TR=8500 milliseconds, TE=90 milliseconds, voxel dimensions = 2 mm isotropic.

The NKI-Rockland high-resolution 3D T1-weighted structural images were obtained using a Magnetization Prepared Rapid Gradient Echo (MPRAGE) sequence with the following parameters: Repetition Time (TR)=1900 milliseconds; Echo Time (TE)=2.52 milliseconds; Inversion Time (TI)=900 milliseconds; flip angle=9 degrees; voxel dimensions=1 mm isotropic (see http://fcon_1000.projects.nitrc.org/indi/enhanced/NKI_MPRAGE.pdf for additional details). Diffusion-Weighted Images (DWI) were acquired with a Diffusion Tensor Imaging (DTI) sequence with 137 diffusion gradient directions with a b-value of 1500 s/mm2. Other relevant parameters include: TR=2400 milliseconds, TE=85 milliseconds, voxel dimensions=2 mm isotropic (see http://fcon_1000.projects.nitrc.org/indi/pro/eNKI_RS_TRT/DIff_137.pdf for additional details).

### 2.1.4 White matter connectome construction

Note that part of the following pipeline is identical to that described in (Bathelt et al., 2019). Diffusion-weighted images were pre-processed to create a brain mask based on the b0-weighted image (FSL BET; Smith, 2002) and to correct for movement and eddy current-induced distortions (eddy; Graham et al., 2016). Subsequently, the diffusion tensor model was fitted and fractional anisotropy (FA) maps were calculated (dtifit). Images with a between-image displacement greater than 3mm as indicated by FSL eddy were excluded from further analysis. All steps were carried out with FSL v5.0.9 and were implemented in a pipeline using NiPyPe v0.13.0 (Gorgolewski et al., 2011). To extract FA values for major white matter tracts, FA images were registered to the FMRIB58 FA template in MNI space using a sequence of rigid, affine, and symmetric diffeomorphic image registration (SyN) as implemented in ANTS v1.9 (Avants et al., 2008). Visual inspection indicated good image registration for all participants. Subsequently, binary masks from a probabilistic white matter atlas (threshold at >50% probability) in the same space were applied to extract FA values for white matter tracts (see below).

Participant movement, particularly in developmental samples, can significantly affect the quality, and, hence, statistical analyses of MRI data. Therefore, we undertook several procedures to ensure adequate MRI data quality and minimize potential biases due to subject movement. First, for the CALM sample, children were trained to lie still inside a realistic mock scanner prior to their actual scans. Secondly, for both samples, all T1-weighted images and FA maps were visually examined by a qualified researcher to remove low quality scans. Lastly, quality of the diffusion-weighted data were evaluated in both samples by calculating the framewise displacement between subsequent volumes in the sequence. Only data with a maximum between-volume displacement below 3mm were included in the analyses. All steps were carried out with FMRIB Software Library v5.0.9 and implemented in the pipeline using NiPyPe v0.13.0 (see https://nipype.readthedocs.io/en/latest/).

### 2.1.5 Neural measures: white matter and fractional anisotropy

To approximate white matter contributions to fluid and crystallized ability, we analyzed fractional anisotropy (FA; see Wandell, 2016). We based our choice of FA on previous studies of white matter in developmental samples (de Mooij et al., 2018; Kievit et al., 2016). We used FA as a general summary metric of white matter microstructure as it cannot directly discern between specific cellular components (e.g. axonal diameter, myelin density, water fraction). Mean FA was computed for 10 bilateral tracts as defined by the Johns Hopkins University DTI-based white matter tractography atlas (see Fig. 1 of Hua et al., 2008): forceps minor (FMin), forceps major (FMaj), anterior thalamic radiations (ATR), cingulate gyrus (CING), superior longitudinal fasciculus (SLF), inferior longitudinal fasciculus (ILF), corticospinal tract (CST), uncinate fasciculus (UNC), cingulum [hippocampus] (CINGh), and inferior fronto-occipital fasciculus (IFOF). Fig. 3 shows the cross-sectional trends of FA across the age range for both samples.

**Fig. 3.**
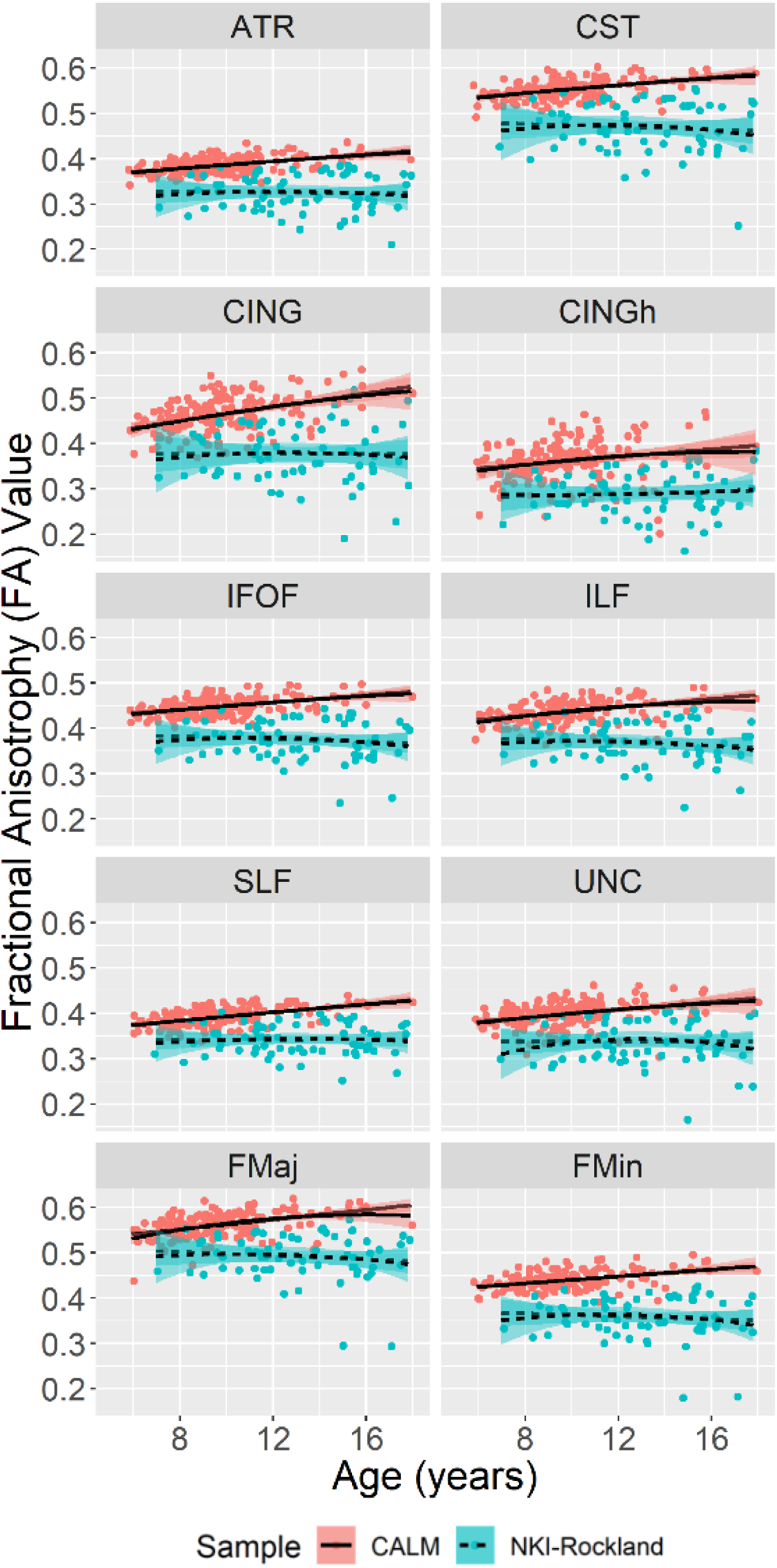
Scatterplots of FA values for all white matter tracts across age for CALM and NKI-Rockland samples. Lines reflect linear fit. Note that the age trends are more pronounced in CALM than in the NKI sample, possibly due to lower sample size in the NKI-Rockland sample (N=65). Lines and shades reflect linear and polynomial fit and 95% confidence intervals, respectively. Solid lines: CALM. Dashed lines: NKI-Rockland. Abbreviations: anterior thalamic radiations (ATR), corticospinal tract (CST), cingulate gyrus (CING), cingulum [hippocampus] (CINGh), inferior fronto-occipital fasciculus (IFOF), inferior longitudinal fasciculus (ILF), superior longitudinal fasciculus (SLF), uncinate fasciculus (UNC), forceps major (FMaj), and forceps minor (FMin).

### 2.1.6 Statistical analyses

We used structural equation modelling (SEM), a multivariate approach that combines latent variables and path modelling to test causal hypotheses (Schreiber et al., 2006) as well as SEM trees, which combine SEM and decision tree paradigms to simultaneously permit exploratory and confirmatory data analysis (Brandmaier et al., 2013).

We performed structural equation modelling (SEM) using the lavaan package version 0.5-22 (Rosseel, 2012) in R (R Core Team, 2018) and versions 2.9.9 and 0.9.12 of the R packages OpenMx (Boker et al., 2011) and semtree (Brandmaier et al., 2013), respectively. To account for missing data and deviations from multivariate normality, we used robust full information maximum likelihood estimator (FIML) with a Yuan-Bentler scaled test statistic (MLR) and robust standard errors (Rosseel, 2012). We evaluated overall model fit via the (Satorra-Bentler scaled) chi-squared test, the comparative fit index (CFI), the standardized root mean squared residuals (SRMR), and the root mean square error of approximation (RMSEA) with its confidence interval (Schermelleh-Engel et al., 2003). Assessment of model fit was defined as: CFI (acceptable fit 0.95-0.97, good fit >0.97), SRMR (acceptable fit 0.05-.10, good fit <0.05), and RMSEA (acceptable fit 0.05-0.08, good fit <0.05). To determine whether gc and gf were separable constructs, we compared a two-factor (gc-gf) model to an single-factor (*g*) model. To investigate if the covariance between gc and gf differed across ages, we conducted multiple group comparisons between younger and older participants based on median splits (CALM split at 8.91 years yielding N=279 young and 272 old; NKI-Rockland split at 11.38 years into N=169 young and N=168 old). Doing so inevitably led to slightly unbalanced numbers of participants with white matter data (CALM: young, N=60 & old, N=105; NKI-Rockland: young, N=19 & old, N=46). To test measurement invariance across age groups (Putnick and Bornstein, 2016), we fit multigroup models (French and Finch, 2008), constraining key parameters across groups. Model comparisons and deviations from measurement invariance were determined using the likelihood ratio test and Akaike information criterion (AIC, see Bozdogan, 1987).

To examine whether white matter tracts made unique contributions to our latent variables we fit Multiple Indicator, Multiple Cause (MIMIC) models (Jöreskog and Goldberger, 1975; Kievit et al., 2012). Lastly, we conducted a SEM tree analysis, a method that combines the confirmatory nature of SEM with the exploratory framework of decision trees (Brandmaier et al., 2013). SEM trees hierarchically and recursively partition datasets based on a covariate (in our case age). This creates data-driven age-groups which show differences in one or more paths of interest. The advantage of SEM trees is that they do not require a-priori decisions as to where potential categorical boundaries between age groups may lie (as was the case in the median split analysis). SEM trees also do not require a-priori knowledge as to the shape of developmental trajectories (as is usually the case when using age as a continuous covariate). Using this technique therefore allowed us to: 1) examine the robustness of findings based on the median age split, and 2) examine whether white matter contributions differed across age groups of younger and older participants in a data-driven way (Hypothesis 4). Therefore, for our SEM tree analyses in CALM and NKI-Rockland, we used age as a continuous covariate. Finally, we used Bonferroni-correction at alpha level .001 to correct for multiple comparisons, and the semtree cross-validation scheme, which “partitions the data for maximizing splits on each variable, then comparing maximum splits across each variable on the rest of the data” (see https://cran.r-project.org/web/packages/semtree/semtree.pdf for more details)

## 3. Results

### 3.1 Covariance among cognitive abilities cannot be captured by a single-factor

In accordance with our <u>preregistered analysis plan</u>, we first describe model fit for the measurement models of the cognitive data only. First, we tested hypothesis 1: that gc and gf are separable constructs in childhood and adolescence. More specifically, we tested the hypothesis that the covariance among scores on cognitive tests would be better captured by a two-factor (gc-gf) model than a single-factor (e.g. *g*) model. In support of this prediction, the single-factor model fit the data poorly: *χ*^2^(27)=317.695, p<.001, CFI=.908, SRMR=.040, RMSEA=.146 [.132 .161], Yuan-Bentler scaling factor=1.090, suggesting that cognitive performance was not well represented by a single-factor. The two-factor (gc-gf) model also displayed poor model fit (*χ*^2^(24)=196.348, p<.001, CFI=.946, SRMR=.046, RMSEA=.119 [.104 .135], Yuan-Bentler scaling factor= 1.087), although it fit significantly better (*χ*^2^Δ=119.41, dfΔ=3, AICΔ=127, p<.001) than the single-factor model.

To investigate the source of poor fit, we examined modification indices (Schermelleh-Engel et al., 2003), which quantify the expected improvement in model fit if a parameter is freed. Modification indices suggested that the Peabody Picture Vocabulary Test had a very strong cross-loading onto the fluid intelligence latent factor. The Peabody Picture Vocabulary Test (PPVT), often considered a crystallized measure in adult populations, asks participants to choose the picture (out of four multiple-choice options) corresponding to the meaning of the word spoken by an examiner. Including a cross-loading between gf and the PPVT drastically improved goodness of fit (*χ*^2^Δ=67.52, dfΔ=1, AICΔ=100, p<.001) to adequate (*χ*^2^(23)=104.533, p<.001, CFI=.975, SRMR=.025, RMSEA=.083 [.067 .099], Yuan-Bentler scaling factor= 1.069). A likely explanation of this result is that such tasks may draw considerably more on executive, gf-like abilities in younger, lower ability samples. For a more thorough investigation of the loading of PPVT across development, see Supplementary Material. Notably, fitting the PPVT as a *solely* fluid task (i.e. removing it as a measurement of gc entirely) did not significantly decrease model fit (*χ*^2^Δ=2.058, dfΔ=1, AICΔ=1, p=.152). Therefore, we decided to proceed with the more parsimonious PPVT gf-only model (*χ*^2^(24)=106.382, p<.001, CFI=.972, SRMR=.025, RMSEA=.082 [.066 .098], Yuan-Bentler scaling factor= 1.073). We note that although this is a data-driven modification, we believe it would likely generalize to samples with similarly low ages and abilities.

Next, we examined whether the single or two-factor model fit best in the NKI-Rockland sample. The single-factor model fit the data adequately (*χ*^2^(14)=41.329, p<.001, CFI=.983, SRMR=.029, RMSEA=.075 [.049 .102], Yuan-Bentler scaling factor=.965). Still, the two-factor model showed considerably better fit (*χ*^2^(12)=19.732, p=.072, CFI= .995, SRMR=.018, RMSEA=.043 [.000 .075], Yuan-Bentler scaling factor=.956) compared to the single-factor model (*χ*^2^Δ=20.661, dfΔ=2, AICΔ=17, p<.001). It should be noted that, given the differences in tasks measured between the samples, gf and working memory were assumed to be measurements of the same latent factor, rather than separable factors. A similar competing model where gf and working memory were modeled as separate constructs with working memory loaded onto gf, similarly to the best-fitting model for the CALM sample (see Fig. 4), showed comparable model fit and converging conclusions with further analyses. Overall, these findings suggested, that for both the CALM and NKI-Rockland samples, a two-factor model with separate gc and gf factors provided a better account of individual differences in intelligence than a single-factor model.

**Fig. 4.**
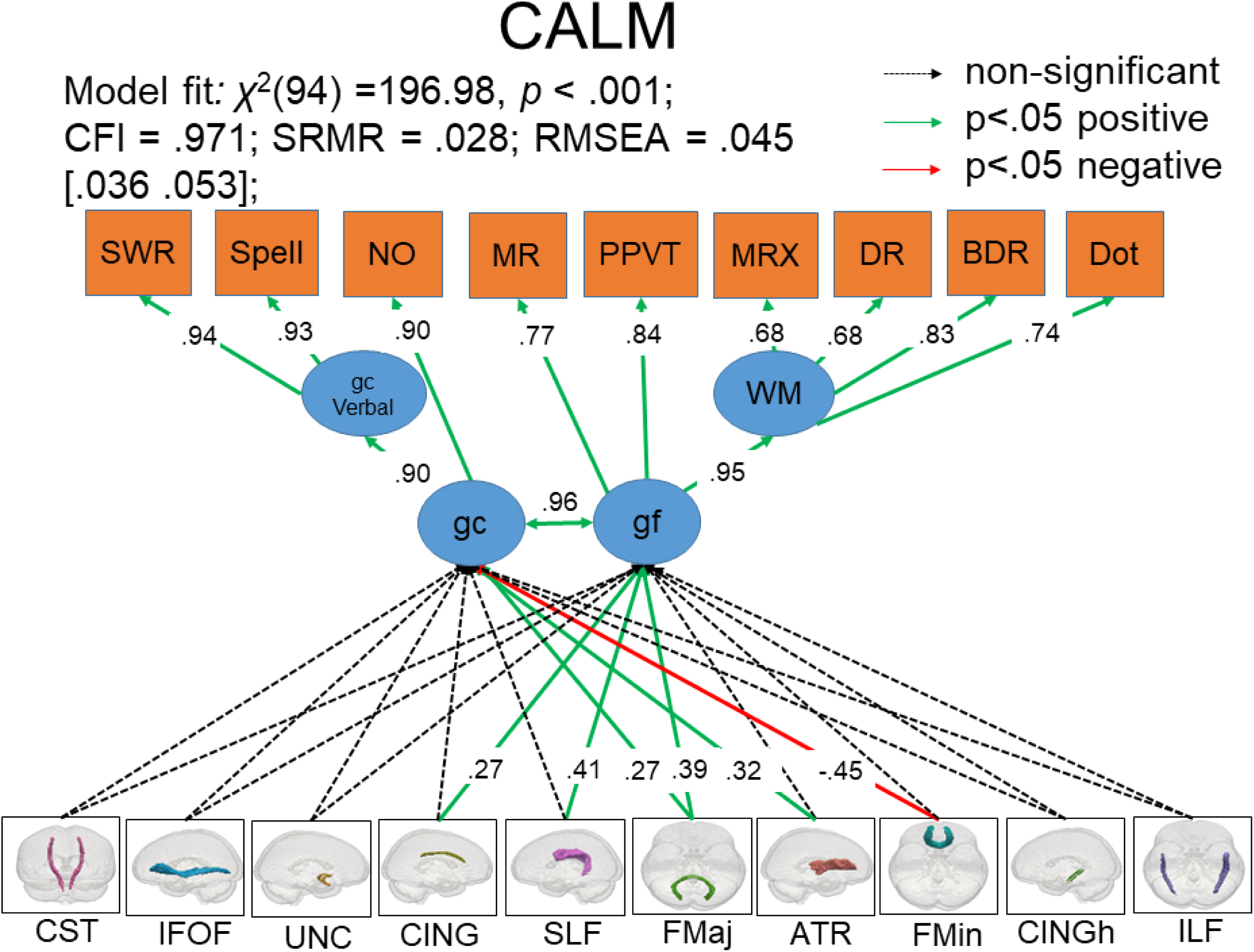
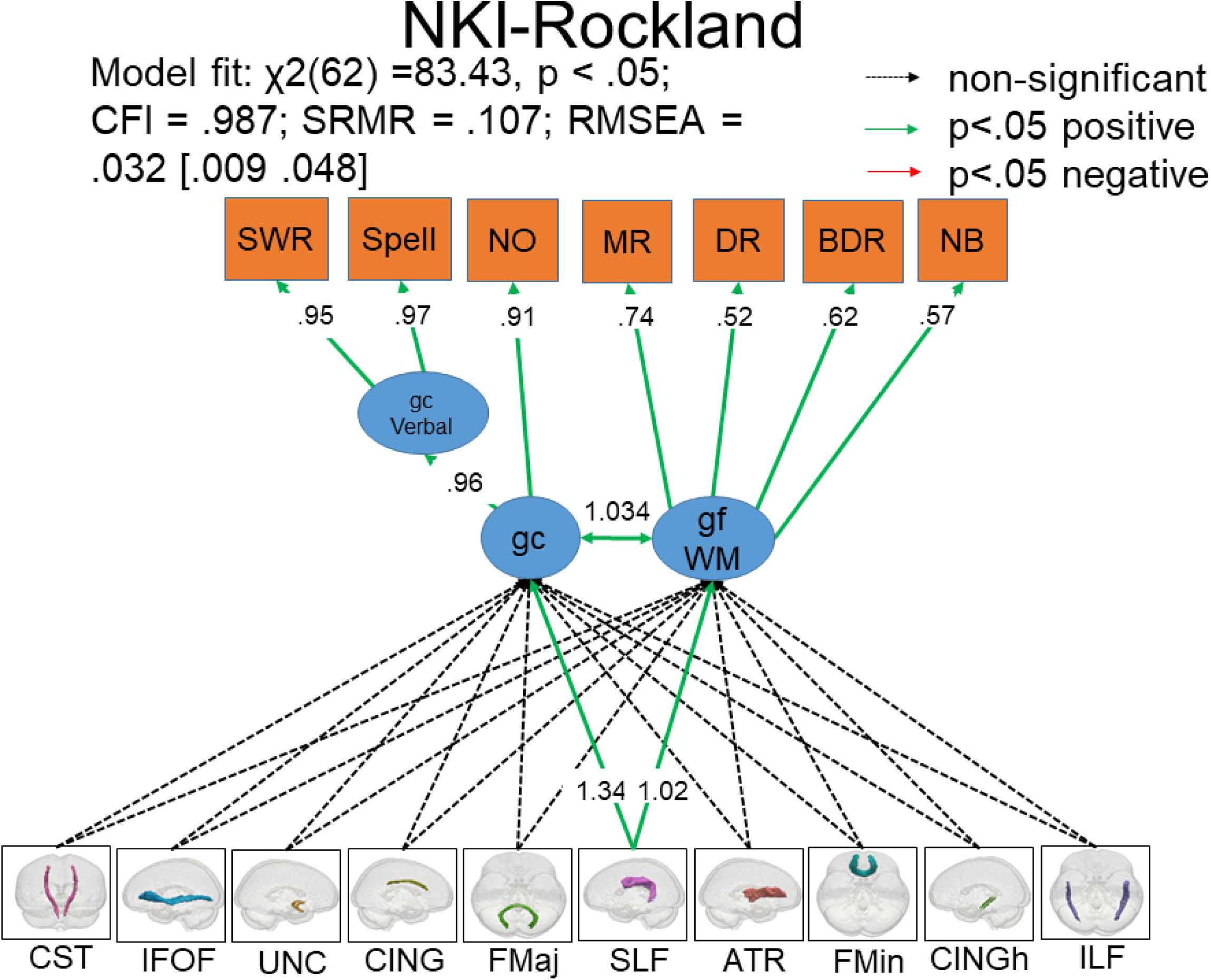
MIMIC models displaying standardized parameter estimates and regression coefficients for all cognitive measures and white matter tracts for complete CALM and NKI-Rockland samples. Dotted, green, and red arrows indicate nonsignificant (>.05), positively significant, and negatively significant path estimates, respectively. Note standardized estimate exceeding 1 in NKI is likely the consequence of highly-correlated factors (Jöreskog, 1999).

### 3.2 Evidence of age differentiation between crystallized and fluid ability

We investigated the relationship between gc and gf in development to see whether we could observe evidence for age differentiation as predicted by hypothesis 2. Age differentiation (e.g. Hülür et al., 2011) would predict decreasing covariance between gc and gf from childhood to adolescence. We fit a multigroup confirmatory factor analysis to assess fit on our younger (N=279) and older (N=272) participant cohorts. The model had acceptable fit (*χ*^2^(48)=142.214, p<.001, CFI= .960, SRMR=.037, RMSEA=.085 [.069 .102], Yuan-Bentler scaling factor= 1.019). However, a likelihood ratio test, showed that model fit did not decrease significantly when imposing equal covariance between gc and gf in the younger and older participant subgroups (*χ*^2^Δ=0.323, dfΔ=1 AICΔ=2, p=.57). This suggested no evidence for age differentiation in the CALM sample. However, the lack of association could be due to limitations of using median splits to investigate age differences when independent (or latent in our case) variables are correlated (Iacobucci et al., 2015). For instance, if the age range of differences in behavioral associations between gc and gf lies elsewhere, the median split may not be sensitive enough to detect it. To test this explicitly, we next fit SEM trees (Brandmaier et al., 2013) to the cognitive data.

We estimated SEM trees in the CALM sample by specifying the cognitive model with age as a continuous covariate. We observed a SEM tree split at age 9.12, yielding two groups (younger participants = 290, older participants = 261). This split was accompanied by a decrease in the unstandardized parameter estimate between gc and gf (from .64 to .59, see Table 3 in section 3.5), providing support for age differentiation using a more exploratory approach (SEM tree: 9.12 versus median split: 8.91). When we fit the two-factor model before and after the SEM tree age split, we found that the correlation between gc and gf increased slightly (from .90 to .92).

**Table 2.** First row: standardized path estimates for cognitive assessments in CALM & NKI-Rockland samples. Second row: raw path estimates with standard errors (parentheses). Third row: 95% confidence intervals [brackets]. NA=not applicable. Note that age groups were determined according to the median split (CALM: 8.91 years, NKI: 11.38 years).

| Relationship | CALM |  | NKI-Rockland |  |
| --- | --- | --- | --- | --- |
|  | Young | Old | Young | Old |
| **gc↔gf(WM)** | 0.89 | 0.93 | 1.01 | 0.81 |
|  | 40.23<br>(5.40) | 44.27 (4.50) | 64.49<br>(8.64) | 12.27<br>(2.68) |
|  | [29.65,<br>50.82] | [ 35.45,<br>53.09] | [47.56,<br>81.43] | [7.02,<br>17.52] |
| **gf→WM** | 0.96 | 0.9 |  |  |
|  | 1.06<br>(.19) | .79 (.09) | NA | NA |
|  | [.69,<br>1.44] | [ .61, .97] |  |  |
| **gf(WM)→MR** | 0.59 | 0.74 | 0.69 | 0.6 |
|  | 1.00<br>(NA) | 1.00 (NA) | 1.00<br>(NA) | 1.00<br>(NA) |
|  | [1.00,<br>1.00] | [1.00, 1.00] | [1.00,<br>1.00] | [1.00,<br>1.00] |
| **gf→PPVT** | 0.75<br>7.49<br>(.84)<br>[5.84,<br>9.14] | 0.76<br>5.45 (.43)<br>[ 4.60, 6.30] | NA | NA |
| **gf(WM)→DR** | 0.56<br>1.00<br>(NA)<br>[1.00,<br>1.00] | 0.68<br>1.00<br>(NA)<br>[1.00, 1.00] | 0.38<br>.12 (.03)<br>[.07, .17] | 0.54<br>.27 (.07)<br>[.13, .40] |
| **gf(WM)→BDR** | 0.76<br>1.01<br>(.12)<br>[.77,<br>1.26] | 0.79<br>.94 (.09)<br>[ .76, 1.12] | 0.5<br>.16 (.03)<br>[.10, .22] | 0.53<br>.30 (.08)<br>[.13, .40] |
| **gfWM→NB** | NA | NA | 0.55<br>.67 (.10)<br>[.48, .87] | 0.35<br>.54 (.14)<br>[.27, .81] |
| **WM→Dot** | 0.58<br>.87 (.12)<br>[.63,<br>1.10] | 0.67<br>1.06 (.12)<br>[ .82, 1.30] | NA | NA |
| **WM→MRX** | 0.59<br>.80 (.11)<br>[.57,<br>1.02] | 0.56<br>.82 (.13)<br>[ .56, 1.08] | NA | NA |
| **gc→ gcV** | 0.89<br>1.00<br>(NA)<br>[1.00,<br>1.00] | 0.79<br>1.00<br>(NA)<br>[1.00, 1.00] | 0.96<br>1.00<br>(NA)<br>[1.00,<br>1.00] | 0.87<br>1.00<br>(NA)<br>[1.00,<br>1.00] |
| **gc→NO** | 0.87<br>.19 (.01)<br>[ .17,<br>.22] | 0.87<br>.54 (.06)<br>[ .43, .65] | 0.9<br>.42 (.03)<br>[.36, .49] | 0.76<br>1.08<br>(.20)<br>[.69,<br>1.48] |
| **gcV→SWR** | 0.94<br>1.00<br>(NA)<br>[1.00,<br>1.00] | 0.91<br>1.00<br>(NA)<br>[1.00, 1.00] | 0.93<br>1.00<br>(NA)<br>[1.00,<br>1.00] | 0.89<br>1.00<br>(NA)<br>[1.00,<br>1.00] |
| **gcV→Spell** | 0.87<br>.28 (.02)<br>[ .25,<br>.31] | 0.91<br>.46 (.03)<br>[ .40, .51] | 0.97<br>.48 (.02)<br>[.44, .52] | 0.88<br>.71 (.06)<br>[.60, .83] |

**Table 3.**
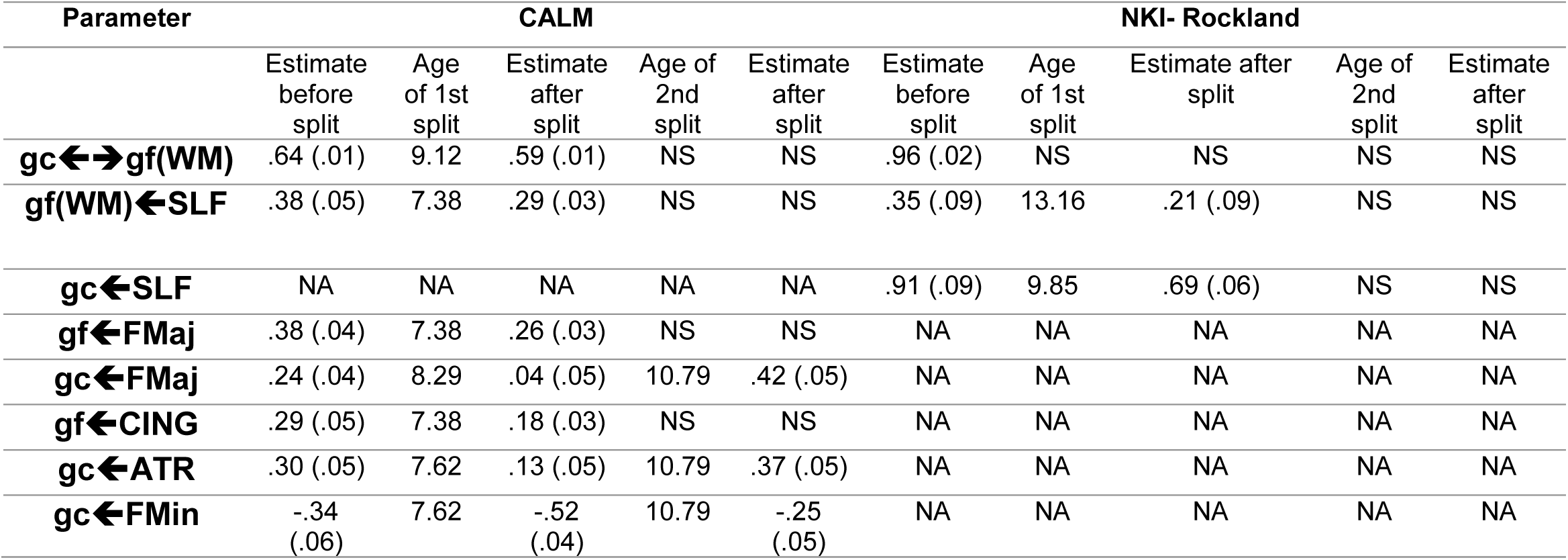
SEM tree Results for CALM & NKI Rockland samples. Note: values listed represent unstandardized estimates and standard errors (parentheses). NS=no SEM tree split, NA=not applicable.

Next, as in the CALM cohort, we fit a multigroup model with younger (N=169) and older (N=168) age groups in the NKI-Rockland sample, which produced good fit (*χ*^2^(24)=33.736, p=.089, CFI=.991, SRMR=.035, RMSEA=.047 [.000 .081], Yuan-Bentler scaling factor=.916). In contrast to CALM, imposing equality constraints on the covariance between gc and gf across age groups revealed a lower gc-gf correlation for the older (.811) compared to the younger participants cohort (1.008). This revealed significant difference in model fit compared to the freely-estimated model (*χ*^2^Δ=61.244, dfΔ=1 AICΔ=46, p<.001). This suggested evidence for age differentiation in the NKI-Rockland sample using multigroup models.

In contrast to the multigroup model outcome, the NKI-Rockland SEM tree model under identical specifications as in CALM failed to produce an age split. A possible explanation is that to penalize for multiple testing we relied on Bonferroni-corrected alpha thresholds for the SEM tree. If, as seems to be the case here, the true split lies (almost) exactly on the median split, then the SEM tree will have slightly less power than conventional multigroup models, as the SEM tree likelihood ratio test is penalized for the number of tests (splits). These differences between analyses methods suggested that the age differentiation observed here is likely modest in size. Taken together, we interpret our findings as evidence for a small, age-specific but suggestive decrease in gc-gf covariance in both cohorts, which is compatible with age differentiation such that, for younger participants, gc and gf factors are almost indistinguishable, whereas for older participants a clearer separation emerges.

### 3.3 Violation of metric invariance suggests differences in relationships among cognitive abilities in childhood and adolescence

Finally, we more closely examined age-related differences in cognitive architecture (e.g. factor loadings) by examining metric invariance (Putnick and Bornstein, 2016). Testing this in the CALM sample as a two-group model by imposing equality constraints on the factor loadings (fully constrained) showed that the freely-estimated model (no factor loading constraints) outperformed the fully-constrained model (*χ*^2^Δ=107.05, dfΔ=7, AICΔ=82, p<.001), indicating that metric invariance was violated. This violation of metric invariance suggested that the relationship between the cognitive tests and latent variables was different in the two age groups. Closer inspection suggested that the differences in loadings were not uniform, but rather showed a more complex pattern of age-related differences (see Table 2 for more details). Some of the most pronounced differences include an increase of the loading of matrix reasoning onto gf as well as increased loading of digit recall and dot matrix onto working memory across age groups.

Similarly, in the NKI-Rockland cohort, the freely-estimated model outperformed the constrained model (*χ*^2^Δ=41.111, dfΔ=5, AICΔ=33, p<.001), indicating that metric invariance was again violated as in CALM. This suggests that the relationship between the cognitive tests and the latent factors differed across age groups. The pattern of factor loadings differed in some respects from CALM. For example, the loading of the N-back task onto gf showed the largest difference across age groups in the NKI-Rockland sample. However, as CALM did not include the N-back task, we cannot directly interpret this as a difference between the cohorts. For detailed comparisons among factor loadings between age groups in both samples, refer to Table 2. The overall pattern in both samples suggested small and varied differences in the relationship between the latent factors and observed scores. A plausible explanation is that the same task draws on a different balance of skills as children differ in age and ability. Our findings concerning the latent factors should be interpreted in this light as it seems likely that in addition to age differentiation (and possibly dedifferentiation) effects, the nature of the factors also differed slightly across the age range studied here.

### 3.4 The neural architecture of gc and gf indicates unique contributions of multiple white matter tracts to cognitive ability

We next focused on the white matter regression coefficients to inspect the neural underpinnings of gc and gf. In line with hypothesis 3, we wanted to explore whether individual white matter tracts made independent contributions to gc and gf. First, we examined whether a single-factor model could account for covariance in white matter microstructure across our ten tracts. If so, then scores on such a latent factor would represent a parsimonious summary for neural integrity. However, this model showed poor fit (*χ*^2^(35)=124.810, p<.001, CFI= .938, SRMR=.039, RMSEA=.132 [.107 .157], Yuan-Bentler scaling factor=1.114), suggesting separate influences from white matter regions in supporting cognitive abilities. To examine whether the white matter tracts showed specific and complementary associations with cognitive performance, we fit a MIMIC model. Doing so, we observed that 5 out of the 10 tracts showed significant relations with gc and/or gf (Fig. 4). Specifically, the anterior thalamic radiations, forceps major, and forceps minor had moderate to strong associations with gc with similar relations seen for gf for the superior longitudinal fasciculus, forceps major, and the cingulate gyrus. Interestingly, the forceps minor exhibited a negative association with gf. This could be due to modeling several highly correlated paths simultaneously since this relationship was not found when only the forceps minor was modeled onto gc (standardized estimate=.426) and gf (standardized estimate=.386, see Tu et al., 2008). Together, individual differences in white matter microstructure explained 32.9% in crystallized and 33.6 % in fluid ability.

As in the CALM sample, the single-factor white matter model produced poor fit (*χ*^2^ (35) =131.637, p<.001, CFI= .924, SRMR=.023, RMSEA=.201 [.165 .238], Yuan-Bentler scaling factor=.950) in the NKI-Rockland sample. Therefore, we fit a multi-tract MIMIC model. The superior longitudinal fasciculus emerged as the only tract to significantly load onto gc or gf (Fig. 4). This result was likely due to lower power associated with a small subset of individuals with white matter data (see Discussion for further investigation). In NKI-Rockland, the same set of tracts explained 29.7% and 26.7% of the variance in gc and gf, respectively, yielding similar joint effect sizes as in the CALM sample. Together, these findings demonstrated generally similar associations between white matter microstructure and cognitive abilities in the CALM and NKI-Rockland samples. Therefore, it seems to be the case that, in both typically and atypically (struggling learners) developing children and adolescents, individual white matter tracts make distinct contributions to crystallized and fluid ability, as more than one tract explains variance in the outcomes (gc and gf) above and beyond all other tracts.

### 3.5 Support for neurocognitive reorganization of crystallized and fluid ability in childhood and adolescence

Lastly, to address our fourth and final preregistered hypothesis, we examined whether brain-behavior associations differed across the developmental age range. We hypothesized that the relationship between the white matter tracts and cognitive abilities would decrease across the age range, in support of the differentiation hypothesis. Using a multigroup model, we compared the strength of brain-behavior relationships between younger and older participants to test whether white matter contributions to gc and gf differed in development. Contrary to our prediction, we observed that, in the CALM sample, a freely estimated model, where the brain-behavior relationships were allowed to vary across age groups, did not outperform the constrained model (*χ*^2^Δ=12.16, dfΔ=10, AICΔ=9, p=.27). This suggested that the contributions of white matter tracts did not vary significantly between age groups when examined using multigroup models.

As before, we estimated a SEM tree model. In contrast to the multigroup model, we observed that multiple white matter tracts *did* differ in their associations with gc and/or gf. These differences manifested in different ways for gc and gf. For example, the correlations between the cingulum, superior longitudinal fasciculus, and forceps major and gf decreased with increasing age, in line with age differentiation. On the other hand, the forceps major, forceps minor and anterior thalamic radiations demonstrated a more complicated pattern with each tract displaying two age splits. For the first split (around age 8), the regression strength decreased before spiking again around age 11 (Table 3, also see Fuhrmann et al., 2019). Given that all first splits showed a decrease between white matter and cognition, and all second splits revealed an increase compared to the first, this suggests a non-monotonic pattern of brain-behavior reorganization that cannot be fully captured by age differentiation or dedifferentiation (Hartung et al., 2018) but may be in line with theories such as Interactive Specialization (Johnson, 2011), which provides a range of mechanisms which may induce age-varying brain-behavior strengths. One hypothesis we have previously offered that may (partially) explain the nature of the age-varying associations between white matter and cognitive performance is the onset of puberty (Fuhrmann et al., 2019, p. 11) and the associated hormonal changes. Previous work has shown that pubertal processes, including differences and changes in hormones such as testosterone, affect diffusion measures in ways that cannot be explained away by (only) age (Menzies et al., 2015). More work in large samples such as ABCD (Volkow et al., 2018), ideally including longitudinal changes in hormone levels, is needed to establish the robustness of this explanation.

Lastly, we performed the same multigroup analysis for the NKI-Rockland MIMIC model, but it failed to converge or produce an age split, likely due to sparsity of the neural data. Therefore, this analysis could not be used to replicate the cutoff age used for multigroup analyses (11.38 years) based on the median split. Further inspection of the only significantly associated tract, the superior longitudinal fasciculus, revealed the same trend for gc and gf with decreased correlations with increasing age (Table 3). Overall, our findings suggest the need for a neurocognitive account of age differentiation-dedifferentiation/reorganization from childhood into adolescence.

## 4. Discussion

### 4.1 Summary of findings

In this <u>preregistered</u> analysis, we examined the cognitive architecture as well as the white matter substrates of fluid and crystallized intelligence in children and adolescents in two developmental samples (CALM and NKI-Rockland). Analyses in both samples indicated that individual differences in intelligence were better captured by two separate but highly correlated factors (gc and gf) of cognitive ability as opposed to a single global factor (*g*). Further analysis suggested that the covariance between these factors decreased slightly from childhood to adolescence, in line with the age differentiation hypothesis of cognitive abilities (Garrett, 1946; Hülür et al., 2011).

We observed multiple, partially independent contributions of specific tracts to individual differences in gc and gf. The clearest associations were observed for the anterior thalamic radiations, cingulum, forceps major, forceps minor, and superior longitudinal fasciculus, all of which have been implicated to play a role in cognitive functioning in childhood and adolescence (Krogsrud et al., 2018; Navas-Sánchez et al., 2014; Peters et al., 2014; Tamnes et al., 2010; Urger et al., 2015; Vollmer et al., 2017). However, except for the superior longitudinal fasciculus, these tracts were not significant in NKI Rockland sample. A possible explanation for this is the difference in imaging sample size between the cohorts (N=165 in the CALM sample versus N=65 in the NKI-Rockland sample). This difference implies sizeable differences in power (73.4% in CALM versus 36.2% in NKI, assuming a standardized effect size of 0.2) to identify weaker individual pathways.

The most consistent association, observed in both samples, was between the superior longitudinal fasciculus, a region known to be important for language and cognition, which significantly contributed to cognitive ability in both CALM (gf only) and NKI-Rockland (gc and gf). The superior longitudinal fasciculus is a long myelinated bidirectional association fiber pathway that runs from anterior to posterior cortical regions and through the major lobes of each hemisphere (Kamali et al., 2014), and has been associated with memory, attention, language, and executive function in childhood and adolescence in both healthy and atypical populations (Frye et al., 2010; Urger et al., 2015). Therefore, given its widespread links throughout the brain, which include temporal and fronto-parietal regions, it is no surprise that it was found to be significantly related to both gc and gf in our samples.

Together, these results are in line with previous research relating fractional anisotropy (FA) and cognitive ability. For instance, Peters et al., 2014 found that age-related differences in cingulum FA mediated differences in executive functioning. Moreover, white matter changes in the forceps major have been linked to higher performance on working memory tasks (Krogsrud et al., 2018). The remaining tracts (superior longitudinal fasciculus and anterior thalamic radiations) have also been positively correlated with verbal and non-verbal cognitive performance in childhood and adolescence (Tamnes et al., 2010; Urger et al., 2015). We also observed more surprising negative pathways, such as between gc and the forceps minor in the CALM sample. However, closer inspection showed that the simple association between forceps minor and gc was *positive*, suggesting the negative pathway is likely the consequence of the simultaneous inclusion of collinear predictors (see Tu et al., 2008).

Finally, using SEM trees (Brandmaier et al., 2013), we observed that white matter contributions to gc and gf differed between participants of different ages. In CALM, the contributions of the cingulum, superior longitudinal fasciculus, and forceps major weakened with increasing age for gf. For gc, however, the forceps major and forceps minor, and the anterior thalamic radiations exhibited a more complex pattern with each tract providing significantly different effects on crystallized intelligence at two distinct time points in development. In NKI-Rockland, the superior longitudinal fasciculus became less associated with both gc and gf. Considering that decreases in white matter relations to gc and gf occurred before covariance decreases found between gc and gf suggest that differences in white matter development may underlie subsequent individual differences in cognition. In a related project (Fuhrmann et al., 2019, Table 6) we observed age-related differences in associations despite focusing on different cognitive factors (processing speed and working memory).

Overall, our findings align with a neurocognitive interpretation of age differentiation-dedifferentiation hypothesis, which would predict that cognitive abilities and their neural substrates become more differentiated (less correlated) until the onset of maturity, followed by an increase (dedifferentiation) in relation to each other until late adulthood (Hartung et al., 2018). However, we note that the evidence for age differentiation-dedifferentiation was not always robust across analyses methods or samples, suggesting only small effect sizes.

### 4.2 Limitations of the present study

First and foremost, all findings here were observed in cross-sectional samples. To better understand effects such as age differentiation and dedifferentiation, future studies will need to model age-related changes within the same individual. The complexity and expense of collecting such longitudinal data has long precluded such investigations, but new cohorts such as the ABCD sample (Volkow et al., 2018) will allow us to model longitudinal changes in the future. Secondly, since the tasks modelled here were not identical between cohorts, detailed interpretations of similarities and differences between the CALM and NKI-Rockland samples should be treated with caution. Therefore, future research comparing cohorts may want to prioritize cohorts with matching tasks to maximize comparability. Thirdly, although the majority of our findings are similar across our cohorts, some differences were observed, particularly in white matter effects. Moreover, although our findings in the SEM tree analysis, of age-related differences in white matter to cognition mapping, are both cross-validated as well as corrected for multiple comparisons, they remain inherently exploratory. Although these findings largely generalize across the two cohorts studied here, further work in larger (such as ABCD, Volkow et al., 2018), more age-heterogeneous (e.g. the Developing Human Connectome Project, Makropoulos et al., 2018) is needed to assess the robustness of these findings. This may reflect statistical variability, differences in sample size and associated differences in power, or true differences between samples. Although the samples here are considerably larger than typical in the field (Poldrack et al., 2017), even larger samples are desirable to gain truly precise estimates of the key parameters, especially regarding measures such as DTI in the NKI-Rockland sample which have a non-trivial proportion of missing data. Moreover, the white matter differences observed could also be due to the scans being obtained at different scanner sites, although this is unlikely to have produced considerable differences for all raw images were processed using the same pipeline, and previous work suggests that FA is quite a robust measure in multi-site comparison (see Vollmar et al., 2010).

In terms of analytical frameworks, here we implement a relatively new analytical framework, called SEM-trees (Brandmaier et al., 2013), to allow for recursive partitioning of our cohorts into age-demarcated subgroups, to capture developmental heterogeneity. SEM-trees have a number of strengths, including considerable flexibility in model specification, implementation in open source software, and the ability to combine measurement and structural model components as well as multiple simultaneous predictors. However, they also have challenges, including potential vulnerability to small fluctuations and overfitting (which may cascade down affecting other partitions), and are certainly not the only choice available to examine model heterogeneity. Alternative analytical strategies, varying in the degree to which they presuppose known group membership or estimate it, include finite mixture models (Zadelaar et al., 2019), Gaussian process structural equation models (e.g. Silva and Gramacy, 2010), latent class and latent profile analysis (Oberski, 2016), general frameworks such as decision trees (McArdle, 2013) and model-based cluster analysis (Fraley and Raftery, 1999), as well as extensions of SEM trees such as SEM forests (Brandmaier et al., 2016). All of these techniques differ in their strengths and weaknesses, ease of implementation, degree of confirmation versus exploration and their flexibility (e.g. can they accommodate latent variables or not). One particularly fruitful avenue for future research is to combine both, using exploratory as well as confirmatory methods to balance discovery and robustness. Here we hopefully illustrate how SEM trees can be one such tool, but would urge the reader to tailor their analytical framework to the question at hand, and be mindful of potential drawbacks. Nonetheless, our view is that SEM trees offer at least one fruitful avenue to formalize hypotheses in developmental cognitive neuroscience which would otherwise often remain mostly verbal.

One concrete concern with SEM trees is the recursive partitioning into subgroups of more modest sample size. Although our two cohorts samples here are considerably larger than typical in the field in terms of their total sample size (Poldrack et al., 2017), the partitioning into subgroups means that some of our parameter estimates are nonetheless based on modest samples, especially regarding measures such as DTI in the NKI-Rockland sample, which have a non-trivial proportion of missing data. Such smaller samples are known to inflate effect sizes (Gelman and Carlin, 2014; Vul et al., 2009). As such, we urge the reader to weigh confidence in the point estimates reported here as a function of sample sizes.

CALM consists of children with referrals for any difficulties related to learning, attention or memory (Holmes et al., 2019). It should be noted that, since CALM is a sample of children and adolescents struggling to learn, and, therefore, ‘atypical’, a large percentage of this cohort had been assigned a diagnosis (36.12%). However, controlling for this possible confound through constrained multigroup models showed this did not affect the results of our models, as was seen in previous work using CALM (Fuhrmann et al., 2019). The NKI-Rockland sample, in contrast, is a United States population representative sample (Nooner et al., 2012). Both samples are composed of large cohorts that underwent extensive phenotyping and population-specific representative sampling. Therefore, we argue that our results generalize to ‘typical’ and ‘atypical’ samples of neurocognitive development.

### 4.3 Conclusions

The present analyses revealed that crystallized and fluid intelligence factors explained a significant amount of variance in test performance in two large child and adolescent samples. These results were found in both typically and atypically (struggling learners) developing cohorts, demonstrating the generalized notion that cognitive ability is better understood as a two-factor rather than a single-factor phenomenon in childhood and adolescence. The addition of white matter microstructure indicated independent contributions from specific white matter tracts known to be involved in cognitive ability. Moreover, further analyses suggested that the associations between neural and behavioral measures differed during development.

Overall, these results support a neurocognitive age differentiation-dedifferentiation hypothesis of cognitive abilities whereby the relation between white matter and cognition become more differentiated (less correlated) in pre-puberty and then dedifferentiate (become more correlated) during early puberty. However, modest subgroup sizes and an inherently exploratory approach such as SEM trees necessitate confirmation in additional, large-scale samples to further quantity the precise developmental differences and changes. Future studies should take this limitation into account when designing experiments attempting to clarify such statements.

## Acknowledgements

The Centre for Attention Learning and Memory (CALM) research clinic is based at and supported by funding from the MRC Cognition and Brain Sciences Unit, University of Cambridge. The Principal Investigators are Joni Holmes (Head of CALM), Susan Gathercole (Chair of CALM Management Committee), Duncan Astle, Tom Manly and Rogier Kievit. Data collection is assisted by a team of researchers and PhD students at the CBU that includes Sarah Bishop, Annie Bryant, Sally Butterfield, Fanchea Daily, Laura Forde, Erin Hawkins, Sinead O’Brien, Cliodhna O’Leary, Joseph Rennie, and Mengya Zhang. The authors wish to thank the many professionals working in children’s services in the South-East and East of England for their support, and to the children and their families for giving up their time to visit the clinic.

We would also like to thank all NKI-Rockland participants and researchers. The NKI-Rockland sample analysed here is the ‘Neurodevelopmental Genomics: Trajectories of Complex Phenotypes Distribution Set 1’ dataset made available through the dbGaP portal under code phs000607.v3.p2.c1. Data was collected using the GOASSESS and Penn Computerized Neurocognitive Battery (CNB) tools, based on the following publications:

Calkins ME, Moore TM, Merikangas KR, et al. The psychosis spectrum in a young U.S. community sample: findings from the Philadelphia Neurodevelopmental Cohort. World Psychiatry. 2014;13(3):296-305. doi:10.1002/wps.20152. https://www.ncbi.nlm.nih.gov/pmc/articles/PMC4219071

Calkins ME, Merikangas KR, Moore TM, et al. The Philadelphia Neurodevelopmental Cohort: constructing a deep phenotyping collaborative. Journal of child psychology and psychiatry, and allied disciplines. 2015;56(12):1356-1369. doi:10.1111/jcpp.12416. https://www.ncbi.nlm.nih.gov/pmc/articles/PMC4598260

Gur RC, Calkins ME, Satterthwaite TD, et al. Neurocognitive Growth Charting in Psychosis Spectrum Youths. JAMA Psychiatry. 2014;71(4):366374. doi:10.1001/jamapsychiatry.2013.4190. https://www.ncbi.nlm.nih.gov/pubmed/24499990 I.L.S.-K. is supported by the Cambridge Trust. D.F., J.B. and G.S. Borgeest are supported by the UK Medical Research Council (MRC). J.A. is funded by the Studienstiftung des deutschen Volkes (German Academic Scholarship Foundation). This project also received funding from the European Union’s Horizon 2020 research and innovation programme (grant agreement number 732592). R. A. Kievit is supported by the Wellcome Trust (Grant No. 107392/Z/15/Z) and the UK Medical Research Council SUAG/014 RG91365.

## Supplementary Material

### Is the Peabody Picture Vocabulary Test a measure of fluid ability?

As a non-preregistered exploratory analysis, we more closely examined the cross-loading of the Peabody Picture Vocabulary Test (PPVT). This task asks participants to select the correct picture (out of four multiple-choice options) corresponding to the meaning of a word spoken by an examiner (Dunn and Dunn, 2007). As discussed previously in the Results section 3.1, modification indices suggested the PPVT should either be cross-loaded or solely loaded onto gf. To better understand this cross-loading, we performed an exploratory (i.e. not part of preregistration) analysis using SEM tree analysis. In this analysis, we allowed the PPVT to load on both gc and gf, and examined whether using age as a covariate yielded a developmental period where the associations between the latent factors and the PPVT task differed. This generated an age split for gf at around age 9.5 whereby the loading of the PPVT decreased (from 1 to .87, unstandardized estimate).

Conversely, for gc the loading remained the same (.12, unstandardized estimate). This suggested the PPVT as commonly implemented behaved as a fluid, rather than a crystallized, task, especially in younger participants of lower ability. Although purportedly a test of crystallized knowledge, the implementation of the PPVT may very well rely on more fluid, executive components including response selection and reasoning, especially in a cohort of children and adolescents with comparatively low overall performance.

A likely explanation for this pattern is that, while PPVT draws on gc, the demanding nature of the task may require more fluid, executive components in younger children, especially in a cohort with comparatively low overall performance (e.g. CALM). Moreover, the surprisingly strong (.83, standardized) association between gf and PPVT in the full sample is similar to previous research in children (Naglieri, 1981) and adults (Bell et al., 2001), although with small, typically developing samples using different statistical methods.

**Supplementary Table 1.** Correlation matrix for cognitive and neural measures in the CALM sample.

| Measure | MR | PPVT | Spell | SWR | NO | DR | Dot | BDR | MRX | ATR | CST | CING | CINGh | IFOF | ILF | SLF | UNC | FMaj | Fmin |
| --- | --- | --- | --- | --- | --- | --- | --- | --- | --- | --- | --- | --- | --- | --- | --- | --- | --- | --- | --- |
| MR |  |  |  |  |  |  |  |  |  |  |  |  |  |  |  |  |  |  |  |
| PPVT | 0.65 |  |  |  |  |  |  |  |  |  |  |  |  |  |  |  |  |  |  |
| Spell | 0.54 | 0.54 |  |  |  |  |  |  |  |  |  |  |  |  |  |  |  |  |  |
| SWR | 0.6 | 0.62 | 0.84 |  |  |  |  |  |  |  |  |  |  |  |  |  |  |  |  |
| NO | 0.69 | 0.63 | 0.74 | 0.69 |  |  |  |  |  |  |  |  |  |  |  |  |  |  |  |
| DR | 0.47 | 0.54 | 0.48 | 0.54 | 0.51 |  |  |  |  |  |  |  |  |  |  |  |  |  |  |
| Dot | 0.56 | 0.56 | 0.49 | 0.49 | 0.6 | 0.45 |  |  |  |  |  |  |  |  |  |  |  |  |  |
| BDR | 0.65 | 0.6 | 0.66 | 0.65 | 0.72 | 0.58 | 0.56 |  |  |  |  |  |  |  |  |  |  |  |  |
| MRX | 0.51 | 0.52 | 0.41 | 0.43 | 0.55 | 0.32 | 0.51 | 0.49 |  |  |  |  |  |  |  |  |  |  |  |
| ATR | 0.29 | 0.34 | 0.43 | 0.45 | 0.38 | 0.2 | 0.24 | 0.31 | 0.16 |  |  |  |  |  |  |  |  |  |  |
| CST | 0.26 | 0.27 | 0.34 | 0.39 | 0.34 | 0.21 | 0.18 | 0.24 | 0.13 | 0.71 |  |  |  |  |  |  |  |  |  |
| CING | 0.33 | 0.42 | 0.43 | 0.42 | 0.44 | 0.32 | 0.32 | 0.36 | 0.23 | 0.69 | 0.62 |  |  |  |  |  |  |  |  |
| CINGh | 0.05 | 0.2 | 0.22 | 0.25 | 0.21 | 0.12 | 0.12 | 0.1 | 0.06 | 0.48 | 0.61 | 0.58 |  |  |  |  |  |  |  |
| IFOF | 0.3 | 0.39 | 0.34 | 0.44 | 0.35 | 0.26 | 0.19 | 0.28 | 0.13 | 0.81 | 0.74 | 0.73 | 0.49 |  |  |  |  |  |  |
| ILF | 0.28 | 0.33 | 0.37 | 0.41 | 0.38 | 0.23 | 0.23 | 0.24 | 0.15 | 0.7 | 0.79 | 0.73 | 0.74 | 0.76 |  |  |  |  |  |
| SLF | 0.4 | 0.4 | 0.4 | 0.44 | 0.43 | 0.29 | 0.31 | 0.33 | 0.2 | 0.75 | 0.8 | 0.72 | 0.54 | 0.8 | 0.79 |  |  |  |  |
| UNC | 0.26 | 0.27 | 0.32 | 0.38 | 0.34 | 0.32 | 0.22 | 0.31 | 0.18 | 0.52 | 0.64 | 0.63 | 0.61 | 0.67 | 0.69 | 0.66 |  |  |  |
| FMaj | 0.39 | 0.38 | 0.36 | 0.4 | 0.43 | 0.31 | 0.26 | 0.36 | 0.2 | 0.7 | 0.74 | 0.65 | 0.59 | 0.69 | 0.74 | 0.73 | 0.61 |  |  |
| Fmin | 0.28 | 0.34 | 0.39 | 0.41 | 0.36 | 0.24 | 0.2 | 0.3 | 0.12 | 0.87 | 0.82 | 0.74 | 0.61 | 0.86 | 0.82 | 0.84 | 0.67 | 0.79 |  |

**Supplementary Table 2.** Correlation matrix for cognitive and neural measures in the NKI-Rockland sample.

| Measure | MR | DR | BDR | SWR | Spell | NO | NB | ATR | CST | CING | CINGh | IFOF | ILF | SLF | UNC | FMaj | Fmin |
| --- | --- | --- | --- | --- | --- | --- | --- | --- | --- | --- | --- | --- | --- | --- | --- | --- | --- |
| MR |  |  |  |  |  |  |  |  |  |  |  |  |  |  |  |  |  |
| DR | 0.16 |  |  |  |  |  |  |  |  |  |  |  |  |  |  |  |  |
| BDR | 0.36 | 0.49 |  |  |  |  |  |  |  |  |  |  |  |  |  |  |  |
| SWR | 0.42 | 0.25 | 0.51 |  |  |  |  |  |  |  |  |  |  |  |  |  |  |
| Spell | 0.34 | 0.16 | 0.43 | 0.89 |  |  |  |  |  |  |  |  |  |  |  |  |  |
| NO | 0.58 | 0.36 | 0.63 | 0.77 | 0.79 |  |  |  |  |  |  |  |  |  |  |  |  |
| NB | 0.25 | 0.23 | 0.42 | 0.36 | 0.17 | 0.41 |  |  |  |  |  |  |  |  |  |  |  |
| ATR | 0.12 | -0.12 | 0.49 | 0.22 | 0.11 | 0.19 | 0.21 |  |  |  |  |  |  |  |  |  |  |
| CST | 0.14 | -0.1 | 0.46 | 0.15 | 0.12 | 0.17 | <.01 | 0.88 |  |  |  |  |  |  |  |  |  |
| CING | 0.17 | -0.1 | 0.36 | 0.08 | -0.05 | 0.06 | 0.44 | 0.59 | 0.4 |  |  |  |  |  |  |  |  |
| CINGh | 0.26 | -0.04 | 0.42 | 0.02 | 0.04 | 0.24 | -0.05 | 0.63 | 0.67 | 0.56 |  |  |  |  |  |  |  |
| IFOF | 0.11 | -0.1 | 0.4 | 0.08 | -0.03 | 0.18 | 0.03 | 0.89 | 0.84 | 0.54 | 0.68 |  |  |  |  |  |  |
| ILF | 0.33 | -0.08 | 0.4 | 0.04 | 0.02 | 0.25 | 0.1 | 0.78 | 0.82 | 0.56 | 0.69 | 0.88 |  |  |  |  |  |
| SLF | 0.02 | -0.06 | 0.4 | 0.22 | 0.18 | 0.26 | 0.13 | 0.83 | 0.79 | 0.53 | 0.5 | 0.87 | 0.84 |  |  |  |  |
| UNC | -0.02 | -0.12 | 0.4 | -0.07 | -0.24 | -0.05 | 0.5 | 0.58 | 0.38 | 0.73 | 0.25 | 0.56 | 0.52 | 0.55 |  |  |  |
| FMaj | 0.08 | -0.11 | 0.35 | 0.04 | 0.03 | 0.21 | 0.07 | 0.74 | 0.82 | 0.23 | 0.46 | 0.73 | 0.73 | 0.73 | 0.39 |  |  |
| Fmin | 0.18 | -0.1 | 0.51 | 0.06 | -0.05 | 0.11 | 0.4 | 0.83 | 0.74 | 0.7 | 0.49 | 0.77 | 0.79 | 0.75 | 0.82 | 0.59 |  |

**Supplementary Fig. 1.**
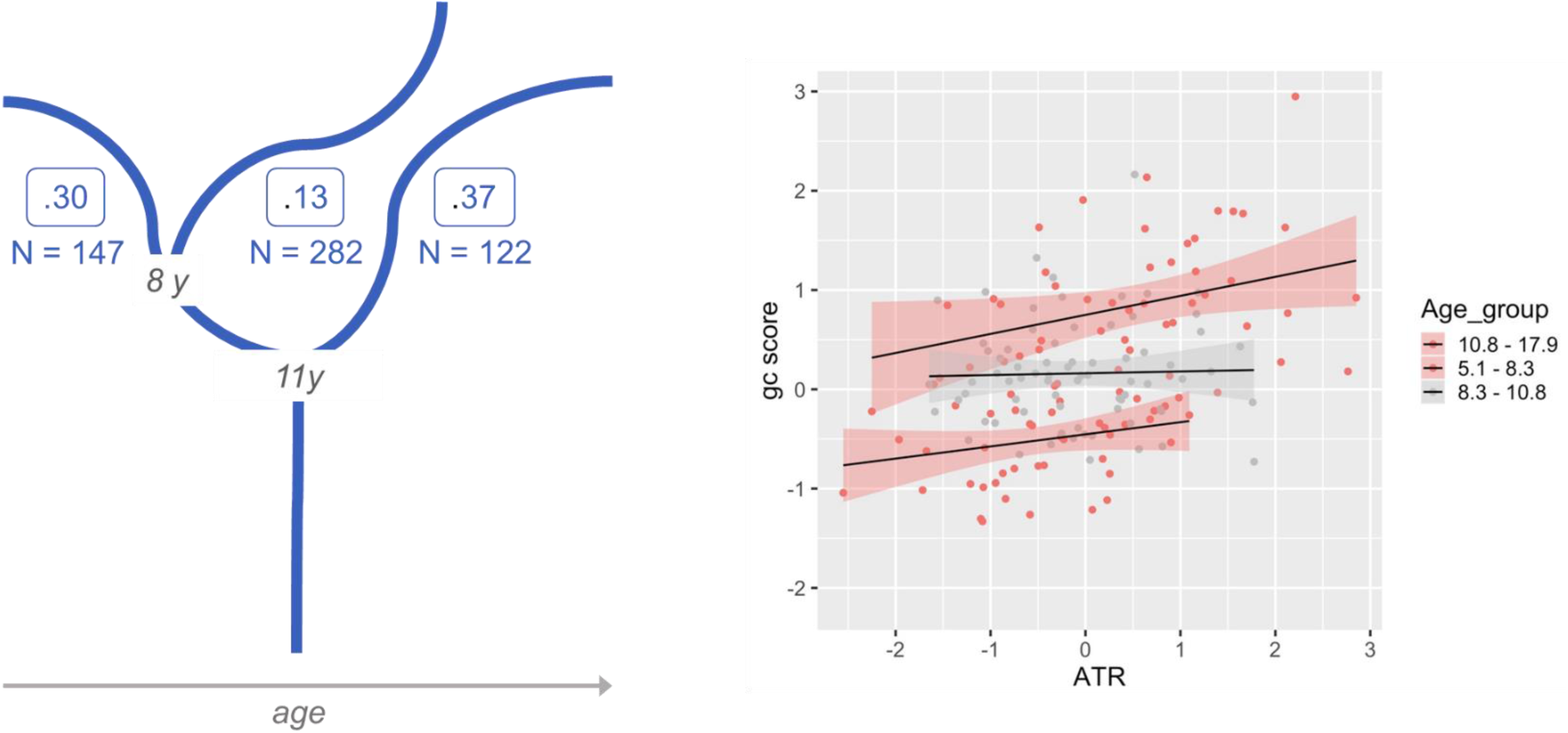
Left: SEM Tree results for the relationship between gc and the anterior thalamic radiations (ATR) (see Table 3 for full results). Figure adapted from Fuhrmann et al., 2019. Right: Visualization the nature of the effect for one path (ATR->gc). The association between FA in the ATR and scores on the gc factor are moderately to strongly positive in the youngest children (r=.22) and the oldest children (r=.29), but effectively absent in the intermediate group (r=0.02). Notably, the reader can see in this figure the steady increase in fractional anisotropy across ages (scaled ATR scores moving rightward) and improvement in gc (gc scores moving upwards).

## Footnotes

1 Gender was coded as either female or male. However, it should be noted that participants might identify themselves as ‘Other’, which, to our knowledge, was not an option according to the biographical protocols used in either sample.

## References

Alloway, T.P., 2007. Automated Working Memory Assessment (AWMA).

Alloway, T.P., Gathercole, S.E., Kirkwood, H., Elliott, J., 2008. Evaluating the validity of the Automated Working Memory Assessment. Educational Psychology 28, 725–734. https://doi.org/10.1080/01443410802243828

Avants, B.B., Epstein, C.L., Grossman, M., Gee, J.C., 2008. Symmetric diffeomorphic image registration with cross-correlation: Evaluating automated labeling of elderly and neurodegenerative brain. Medical Image Analysis, Special Issue on The Third International Workshop on Biomedical Image Registration – WBIR 2006 12, 26–41. https://doi.org/10.1016/j.media.2007.06.004

Bathelt, J., Johnson, A., Zhang, M., Astle, D.E., 2019. The cingulum as a marker of individual differences in neurocognitive development. Scientific Reports 9, 2281. https://doi.org/10.1038/s41598-019-38894-z

Bell, N.L., Lassiter, K.S., Matthews, T.D., Hutchinson, M.B., 2001. Comparison of the Peabody Picture Vocabulary Test—Third Edition and Wechsler Adult Intelligence Scale—Third Edition with university students. Journal of Clinical Psychology 57, 417–422. https://doi.org/10.1002/jclp.1024

Bickley, P.G., Keith, T.Z., Wolfle, L.M., 1995. The three-stratum theory of cognitive abilities: Test of the structure of intelligence across the life span. Intelligence 20, 309–328. https://doi.org/10.1016/0160-2896(95)90013-6

Boker, S., Neale, M., Maes, H., Wilde, M., Spiegel, M., Brick, T., Spies, J., Estabrook, R., Kenny, S., Bates, T., Mehta, P., Fox, J., 2011. OpenMx: An Open Source Extended Structural Equation Modeling Framework. Psychometrika 76, 306–317. https://doi.org/10.1007/s11336-010-9200-6

Bozdogan, H., 1987. Model selection and Akaike’s Information Criterion (AIC): The general theory and its analytical extensions. Psychometrika 52, 345–370. https://doi.org/10.1007/BF02294361

Brandmaier, A.M., Prindle, J.J., McArdle, J.J., Lindenberger, U., 2016. Theory-guided exploration with structural equation model forests. Psychological Methods 21, 566–582. https://doi.org/10.1037/met0000090

Brandmaier, A.M., von Oertzen, T., McArdle, J.J., Lindenberger, U., 2013. Structural equation model trees. Psychological Methods 18, 71–86. https://doi.org/10.1037/a0030001

Calvin, C.M., Deary, I.J., Fenton, C., Roberts, B.A., Der, G., Leckenby, N., Batty, G.D., 2011. Intelligence in youth and all-cause-mortality: systematic review with meta-analysis. International Journal of Epidemiology 40, 626–644. https://doi.org/10.1093/ije/dyq190

Cattell, R.B., 1967. The theory of fluid and crystallized general intelligence checked at the 5-6 year-old level. British Journal of Educational Psychology 37, 209–224. https://doi.org/10.1111/j.2044-8279.1967.tb01930.x

de Mooij, S.M.M., Henson, R.N.A., Waldorp, L.J., Kievit, R.A., 2018. Age Differentiation within Gray Matter, White Matter, and between Memory and White Matter in an Adult Life Span Cohort. The Journal of Neuroscience 38, 5826–5836. https://doi.org/10.1523/JNEUROSCI.1627-17.2018

Deary, I.J., Penke, L., Johnson, W., 2010. The neuroscience of human intelligence differences. Nature Reviews Neuroscience 11, 201–211. https://doi.org/10.1038/nrn2793

Deary, I.J., Strand, S., Smith, P., Fernandes, C., 2007. Intelligence and educational achievement. Intelligence 35, 13–21. https://doi.org/10.1016/j.intell.2006.02.001

Dunn, L.M., Dunn, D.M., 2007. PPVT-4: Peabody picture vocabulary test.

Fraley, C., Raftery, A.E., 1999. MCLUST: Software for Model-Based Cluster and Discriminant Analysis. Journal of Classification 16, 297–306.

French, B.F., Finch, W.H., 2008. Multigroup Confirmatory Factor Analysis: Locating the Invariant Referent Sets. Structural Equation Modeling: A Multidisciplinary Journal 15, 96–113. https://doi.org/10.1080/10705510701758349

Frye, R.E., Hasan, K., Malmberg, B., Desouza, L., Swank, P., Smith, K., Landry, S., 2010. Superior longitudinal fasciculus and cognitive dysfunction in adolescents born preterm and at term: Superior Longitudinal Fasciculus and Cognitive Deficits. Developmental Medicine & Child Neurology 52, 760–766. https://doi.org/10.1111/j.1469-8749.2010.03633.x

Fuhrmann, D., Simpson-Kent, I.L., Bathelt, J., Kievit, R.A., 2019. A Hierarchical Watershed Model of Fluid Intelligence in Childhood and Adolescence. Cerebral Cortex 1–14.

Garrett, H.E., 1946. A developmental theory of intelligence. The American Psychologist 1, 372–378. http://dx.doi.org.ezp.lib.cam.ac.uk/10.1037/h0056380

Gelman, A., Carlin, J., 2014. Beyond Power Calculations: Assessing Type S (Sign) and Type M (Magnitude) Errors. Perspect Psychol Sci 9, 641–651. https://doi.org/10.1177/1745691614551642

Gignac, G.E., 2014. Dynamic mutualism versus g factor theory: An empirical test. Intelligence 42, 89–97. https://doi.org/10.1016/j.intell.2013.11.004

Gorgolewski, K., Burns, C.D., Madison, C., Clark, D., Halchenko, Y.O., Waskom, M.L., Ghosh, S.S., 2011. Nipype: A Flexible, Lightweight and Extensible Neuroimaging Data Processing Framework in Python. Front. Neuroinform. 5. https://doi.org/10.3389/fninf.2011.00013

Graham, M.S., Drobnjak, I., Zhang, H., 2016. Realistic simulation of artefacts in diffusion MRI for validating post-processing correction techniques. NeuroImage 125, 1079–1094. https://doi.org/10.1016/j.neuroimage.2015.11.006

Gur, R.C., Richard, J., Hughett, P., Calkins, M.E., Macy, L., Bilker, W.B., Brensinger, C., Gur, R.E., 2010. A cognitive neuroscience-based computerized battery for efficient measurement of individual differences: Standardization and initial construct validation. Journal of Neuroscience Methods 187, 254–262. https://doi.org/10.1016/j.jneumeth.2009.11.017

Hartung, J., Doebler, P., Schroeders, U., Wilhelm, O., 2018. Dedifferentiation and differentiation of intelligence in adults across age and years of education. Intelligence 69, 37–49. https://doi.org/10.1016/j.intell.2018.04.003

Holmes, J., Bryant, A., Gathercole, S.E., 2018. A transdiagnostic study of children with problems of attention, learning and memory (CALM). bioRxiv. https://doi.org/10.1101/303826

Holmes, J., Bryant, A., Gathercole, S.E., the CALM Team, 2019. Protocol for a transdiagnostic study of children with problems of attention, learning and memory (CALM). BMC Pediatrics 19. https://doi.org/10.1186/s12887-018-1385-3

Horn, J.L., Cattell, R.B., 1967. Age differences in fluid and crystallized intelligence. Acta Psychologica 26, 107–129. https://doi.org/10.1016/0001-6918(67)90011-X

Hua, K., Zhang, J., Wakana, S., Jiang, H., Li, X., Reich, D.S., Calabresi, P.A., Pekar, J.J., van Zijl, P.C.M., Mori, S., 2008. Tract probability maps in stereotaxic spaces: Analyses of white matter anatomy and tract-specific quantification. NeuroImage 39, 336–347. https://doi.org/10.1016/j.neuroimage.2007.07.053

Hülür, G., Wilhelm, O., Robitzsch, A., 2011. Intelligence Differentiation in Early Childhood. Journal of Individual Differences 32, 170–179. https://doi.org/10.1027/1614-0001/a000049

Iacobucci, D., Posavac, S.S., Kardes, F.R., Schneider, M.J., Popovich, D.L., 2015. The median split: Robust, refined, and revived. Journal of Consumer Psychology 25, 690–704. https://doi.org/10.1016/j.jcps.2015.06.014

Johnson, M.H., 2011. Interactive Specialization: A domain-general framework for human functional brain development? Developmental Cognitive Neuroscience 1, 7–21. https://doi.org/10.1016/j.dcn.2010.07.003

Jöreskog, K.G., Goldberger, A.S., 1975. Estimation of a Model with Multiple Indicators and Multiple Causes of a Single Latent Variable. Journal of the American Statistical Association 70, 631–639.

Juan-Espinosa, M., García, L.F., Colom, R., Abad, F.J., 2000. Testing the age related differentiation hypothesis through the Wechsler’s scales. Personality and Individual Differences 29, 1069–1075. https://doi.org/10.1016/S0191-8869(99)00254-8

Kamali, A., Sair, H.I., Radmanesh, A., Hasan, K.M., 2014. Decoding the superior parietal lobule connections of the superior longitudinal fasciculus/arcuate fasciculus in the human brain. Neuroscience 277, 577–583. https://doi.org/10.1016/j.neuroscience.2014.07.035

Kaufman, A.S., 1975. Factor analysis of the WISC-R at 11 age levels between 6 1/2 and 16 1/2 years. Journal of Consulting and Clinical Psychology 43, 135–147.

Kievit, R.A., Davis, S.W., Griffiths, J., Correia, M.M., Cam-CAN Henson, R.N., 2016. A watershed model of individual differences in fluid intelligence. Neuropsychologia 91, 186–198. https://doi.org/10.1016/j.neuropsychologia.2016.08.008

Kievit, R.A., van Rooijen, H., Wicherts, J.M., Waldorp, L.J., Kan, K.-J., Scholte, H.S., Borsboom, D., 2012. Intelligence and the brain: A model-based approach. Cognitive Neuroscience 3, 89–97. https://doi.org/10.1080/17588928.2011.628383

Koenis, M.M.G., Brouwer, R.M., Swagerman, S.C., Soelen, I.L.C. van, Boomsma, D.I., Pol, H.E.H., 2018. Association between structural brain network efficiency and intelligence increases during adolescence. Human Brain Mapping 39, 822–836. https://doi.org/10.1002/hbm.23885

Koenis, M.M.G., Brouwer, R.M., van den Heuvel, M.P., Mandl, R.C.W., van Soelen, I.L.C., Kahn, R.S., Boomsma, D.I., Hulshoff Pol, H.E., 2015. Development of the brain’s structural network efficiency in early adolescence: A longitudinal DTI twin study. Hum. Brain Mapp. 36, 4938–4953. https://doi.org/10.1002/hbm.22988

Krogsrud, S.K., Fjell, A.M., Tamnes, C.K., Grydeland, H., Due-Tønnessen, P., Bjørnerud, A., Sampaio-Baptista, C., Andersson, J., Johansen-Berg, H., Walhovd, K.B., 2018. Development of white matter microstructure in relation to verbal and visuospatial working memory—A longitudinal study. PLOS ONE 13, e0195540. https://doi.org/10.1371/journal.pone.0195540

Makropoulos, A., Robinson, E.C., Schuh, A., Wright, R., Fitzgibbon, S., Bozek, J., Counsell, S.J., Steinweg, J., Vecchiato, K., Passerat-Palmbach, J., Lenz, G., Mortari, F., Tenev, T., Duff, E.P., Bastiani, M., Cordero-Grande, L., Hughes, E., Tusor, N., Tournier, J.-D., Hutter, J., Price, A.N., Teixeira, R.P.A.G., Murgasova, M., Victor, S., Kelly, C., Rutherford, M.A., Smith, S.M., Edwards, A.D., Hajnal, J.V., Jenkinson, M., Rueckert, D., 2018. The developing human connectome project: A minimal processing pipeline for neonatal cortical surface reconstruction. NeuroImage 173, 88–112. https://doi.org/10.1016/j.neuroimage.2018.01.054

McArdle, J.J., 2013. Exploratory Data Mining Using Decision Trees in the Behavioral Sciences [WWW Document]. Contemporary Issues in Exploratory Data Mining in the Behavioral Sciences. https://doi.org/10.4324/9780203403020-10

McArdle, J.J., Hamagami, F., Meredith, W., Bradway, K.P., 2000. Modeling the dynamic hypotheses of Gf–Gc theory using longitudinal life-span data. Learning and Individual Differences 12, 53–79. https://doi.org/10.1016/S1041-6080(00)00036-4

Menzies, L., Goddings, A.-L., Whitaker, K.J., Blakemore, S.-J., Viner, R.M., 2015. The effects of puberty on white matter development in boys. Developmental Cognitive Neuroscience, Proceedings from the inaugural Flux Congress; towards an integrative developmental cognitive neuroscience 11, 116–128. https://doi.org/10.1016/j.dcn.2014.10.002

Naglieri, J.A., 1981. Concurrent validity of the revised Peabody Picture Vocabulary Test. Psychology in the Schools 18, 286–289. https://doi.org/10.1002/1520-6807(198107)18:3<286::AID-PITS2310180306>3.0.CO;2-1

Navas-Sánchez, F.J., Alemán-Gómez, Y., Sánchez-Gonzalez, J., Guzmán-De-Villoria, J.A., Franco, C., Robles, O., Arango, C., Desco, M., 2014. White matter microstructure correlates of mathematical giftedness and intelligence quotient: White Matter Microstructure. Human Brain Mapping 35, 2619–2631. https://doi.org/10.1002/hbm.22355

Nooner, K.B., Colcombe, S.J., Tobe, R.H., Mennes, M., Benedict, M.M., Moreno, A.L., Panek, L.J., Brown, S., Zavitz, S.T., Li, Q., Sikka, S., Gutman, D., Bangaru, S., Schlachter, R.T., Kamiel, S.M., Anwar, A.R., Hinz, C.M., Kaplan, M.S., Rachlin, A.B., Adelsberg, S., Cheung, B., Khanuja, R., Yan, C., Craddock, C.C., Calhoun, V., Courtney, W., King, M., Wood, D., Cox, C.L., Kelly, A.M.C., Di Martino, A., Petkova, E., Reiss, P.T., Duan, N., Thomsen, D., Biswal, B., Coffey, B., Hoptman, M.J., Javitt, D.C., Pomara, N., Sidtis, J.J., Koplewicz, H.S., Castellanos, F.X., Leventhal, B.L., Milham, M.P., 2012. The NKI-Rockland Sample: A Model for Accelerating the Pace of Discovery Science in Psychiatry. Frontiers in Neuroscience 6. https://doi.org/10.3389/fnins.2012.00152

Oberski, D., 2016. Mixture Models: Latent Profile and Latent Class Analysis, in: Robertson, J., Kaptein, M. (Eds.), Modern Statistical Methods for HCI, Human–Computer Interaction Series. Springer International Publishing, Cham, pp. 275–287. https://doi.org/10.1007/978-3-319-26633-6_12

Peters, B.D., Ikuta, T., DeRosse, P., John, M., Burdick, K.E., Gruner, P., Prendergast, D.M., Szeszko, P.R., Malhotra, A.K., 2014. Age-Related Differences in White Matter Tract Microstructure Are Associated with Cognitive Performance from Childhood to Adulthood. Biological Psychiatry 75, 248–256. https://doi.org/10.1016/j.biopsych.2013.05.020

Poldrack, R.A., Baker, C.I., Durnez, J., Gorgolewski, K.J., Matthews, P.M., Munafò, M.R., Nichols, T.E., Poline, J.-B., Vul, E., Yarkoni, T., 2017. Scanning the horizon: towards transparent and reproducible neuroimaging research. Nature Reviews Neuroscience 18, 115–126. https://doi.org/10.1038/nrn.2016.167

Putnick, D.L., Bornstein, M.H., 2016. Measurement invariance conventions and reporting: The state of the art and future directions for psychological research. Developmental Review 41, 71–90. https://doi.org/10.1016/j.dr.2016.06.004

R Core Team, 2018. R: A Language and Environment for Statistical Computing. R Foundation for Statistical Computing, Vienna.

Rosseel, Y., 2012. lavaan: An R Package for Structural Equation Modeling. Journal of Statistical Software 48. https://doi.org/10.18637/jss.v048.i02

Schaie, K.W., 1994. The course of adult intellectual development. American Psychologist 49, 304–313. https://doi.org/10.1037//0003-066X.49.4.304

Schermelleh-Engel, K., Moosbrugger, H., Müller, H., 2003a. Evaluating the Fit of Structural Equation Models: Tests of Significance and Descriptive Goodness-of Fit Measures. Methods of Psychological Research 23–74.

Schermelleh-Engel, K., Moosbrugger, H., Müller, H., 2003b. Evaluating the Fit of Structural Equation Models: Tests of Significance and Descriptive Goodness-of-Fit Measures 8, 52.

Schneider, W.J., McGrew, K.S., 2012. The Cattell-Horn-Carroll model of intelligence, in: Contemporary Intellectual Assessment: Theories, Tests, and Issues, 3rd Ed. The Guilford Press, New York, NY, US, pp. 99–144.

Schreiber, J.B., Nora, A., Stage, F.K., Barlow, E.A., King, J., 2006. Reporting Structural Equation Modeling and Confirmatory Factor Analysis Results: A Review. The Journal of Educational Research 99, 323–338. https://doi.org/10.3200/JOER.99.6.323-338

Silva, R., Gramacy, R.B., 2010. Gaussian Process Structural Equation Models with Latent Variables. 1002.4802 [cs, stat].

Smith, S.M., 2002. Fast robust automated brain extraction. Human Brain Mapping 17, 143–155. https://doi.org/10.1002/hbm.10062

Spearman, C., 1904. “General Intelligence,” Objectively Determined and Measured. The American Journal of Psychology 15, 201–292.

Tamnes, C.K., Østby, Y., Walhovd, K.B., Westlye, L.T., Due-Tønnessen, P., Fjell, A.M., 2010. Intellectual abilities and white matter microstructure in development: A diffusion tensor imaging study. Human Brain Mapping 31, 1609–1625. https://doi.org/10.1002/hbm.20962

Tideman, E., Gustafsson, J.-E., 2004. Age-related differentiation of cognitive abilities in ages 3–7. Personality and Individual Differences 36, 1965–1974. https://doi.org/10.1016/j.paid.2003.09.004

Tu, Y.-K., Gunnell, D., Gilthorpe, M.S., 2008. Simpson’s Paradox, Lord’s Paradox, and Suppression Effects are the same phenomenon – the reversal paradox. Emerg Themes Epidemiol 5, 2. https://doi.org/10.1186/1742-7622-5-2

Urger, S.E., De Bellis, M.D., Hooper, S.R., Woolley, D.P., Chen, S.D., Provenzale, J., 2015. The Superior Longitudinal Fasciculus in Typically Developing Children and Adolescents: Diffusion Tensor Imaging and Neuropsychological Correlates. Journal of Child Neurology 30, 9–20. https://doi.org/10.1177/0883073813520503

Volkow, N.D., Koob, G.F., Croyle, R.T., Bianchi, D.W., Gordon, J.A., Koroshetz, W.J., Pérez-Stable, E.J., Riley, W.T., Bloch, M.H., Conway, K., Deeds, B.G., Dowling, G.J., Grant, S., Howlett, K.D., Matochik, J.A., Morgan, G.D., Murray, M.M., Noronha, A., Spong, C.Y., Wargo, E.M., Warren, K.R., Weiss, S.R.B., 2018. The conception of the ABCD study: From substance use to a broad NIH collaboration. Developmental Cognitive Neuroscience 32, 4–7. https://doi.org/10.1016/j.dcn.2017.10.002

Vollmar, C., O’Muircheartaigh, J., Barker, G.J., Symms, M.R., Thompson, P., Kumari, V., Duncan, J.S., Richardson, M.P., Koepp, M.J., 2010. Identical, but not the same: Intra-site and inter-site reproducibility of fractional anisotropy measures on two 3.0T scanners. NeuroImage 51, 1384–1394. https://doi.org/10.1016/j.neuroimage.2010.03.046

Vollmer, B., Lundequist, A., Mårtensson, G., Nagy, Z., Lagercrantz, H., Smedler, A.-C., Forssberg, H., 2017. Correlation between white matter microstructure and executive functions suggests early developmental influence on long fibre tracts in preterm born adolescents. PLOS ONE 12, e0178893. https://doi.org/10.1371/journal.pone.0178893

Vul, E., Harris, C., Winkielman, P., Pashler, H., 2009. Puzzlingly High Correlations in fMRI Studies of Emotion, Personality, and Social Cognition. Perspect Psychol Sci 4, 274–290. https://doi.org/10.1111/j.1745-6924.2009.01125.x

Wandell, B.A., 2016. Clarifying Human White Matter. Annual Review of Neuroscience 39, 103–128. https://doi.org/10.1146/annurev-neuro-070815-013815

Wechsler, D., 2011. Wechsler Abbreviated Scales of Intelligence-Second Edition (WASI-II).

Wechsler, D., 2005. Wechsler Individual Achievement Test-Second UK Edition (WIAT-II).

Wechsler, D., 1999. Wechsler Abbreviated Scales of Intelligence.

Westerhausen, R., Friesen, C.-M., Rohani, D.A., Krogsrud, S.K., Tamnes, C.K., Skranes, J.S., Håberg, A.K., Fjell, A.M., Walhovd, K.B., 2018. The corpus callosum as anatomical marker of intelligence? A critical examination in a large-scale developmental study. Brain Structure and Function 223, 285–296. https://doi.org/10.1007/s00429-017-1493-0

Zadelaar, J.N., Weeda, W.D., Waldorp, L.J., Van Duijvenvoorde, A.C.K., Blankenstein, N.E., Huizenga, H.M., 2019. Are individual differences quantitative or qualitative? An integrated behavioral and fMRI MIMIC approach. NeuroImage 202, 116058. https://doi.org/10.1016/j.neuroimage.2019.116058

